# The localization of PHRAGMOPLAST ORIENTING KINESIN1 at the division site depends on two microtubule binding proteins TANGLED1 and AUXIN-INDUCED-IN-ROOT-CULTURES9 in Arabidopsis

**DOI:** 10.1101/2022.04.27.489732

**Authors:** Alison M. Mills, Victoria H. Morris, Carolyn G Rasmussen

## Abstract

Proper plant growth and development requires spatial coordination of cell divisions. Two unrelated microtubule-binding proteins, TANGLED1 (TAN1) and AUXIN-INDUCED-IN-ROOT-CULTURES9 (AIR9), are together required for normal growth and division-plane orientation in Arabidopsis. *tan1 air9* double mutants have synthetic growth and division-plane orientation defects while single mutants lack obvious defects. Here we show that the division-site localized protein, PHRAGMOPLAST-ORIENTING-KINESIN1 (POK1), was aberrantly lost from the division site during metaphase and telophase in *tan1 air9* mutants. Since TAN1 and POK1 interact via the first 132 amino acids of TAN1 (TAN1_1-132_), we assessed its localization and function in the *tan1 air9* double mutant. TAN1_1-132_ rescued *tan1 air9* mutant phenotypes and localized to the division site in telophase. However, replacing six amino-acid residues within TAN1_1-132_ that disrupts POK1-TAN1 interaction in the yeast-two-hybrid system caused loss of both rescue and division-site localization of TAN1_1-132_ in *tan1 air9* mutants. Full-length TAN1 with the same alanine substitutions had defects in phragmoplast guidance and reduced TAN1 and POK1 localization at the division site but rescued most *tan1 air9* mutant phenotypes. Together, these data suggest that TAN1 and AIR9 are required for POK1 localization, and yet unknown proteins may stabilize TAN1-POK1 interactions.

**One sentence summary:** Specific amino acids within TAN1 are required for its correct localization and function partially through interaction with POK1; both TAN1 and AIR9 mediate POK1 division site localization.

## Introduction

Division plane orientation is important for many aspects of plant, microbial, and animal development, particularly growth and patterning. Division plane orientation is especially relevant for plant cells which are encased in cell walls, and unable to migrate (Rasmussen and Bellinger, 2018; Livanos and Müller, 2019; Facette et al., 2018; Wu et al., 2018). Positioning and construction of the new cell wall (cell plate) during cytokinesis involves two microtubule- and microfilament-rich cytoskeletal structures, the preprophase band (PPB) and the phragmoplast respectively (Smertenko et al., 2017). The PPB is a ring of microtubules, microfilaments, and proteins that forms at the cell cortex just beneath the plasma membrane during G2: this region is defined as the cortical division zone (Van Damme, 2009; Smertenko et al., 2017; Li et al., 2015). The cortical division zone is characterized by active endocytosis mediated by TPLATE-clathrin coated vesicles that may deplete actin and the actin-binding kinesin like-protein KCA1/KAC1 (Vanstraelen et al., 2004; Suetsugu et al., 2010; Karahara et al., 2009; Kojo et al., 2013; Panteris, 2008; Hoshino et al., 2003). After nuclear envelope breakdown, the PPB disassembles and the metaphase spindle, an antiparallel microtubule array with its plus-ends directed toward the middle of the cell, forms (Dixit and Cyr, 2002). After the chromosomes are separated, the phragmoplast is constructed from spindle remnants to form another antiparallel array of microtubules (Lee and Liu, 2019). The phragmoplast microtubules are tracks for the movement of vesicles containing cell wall materials towards the forming cell plate (McMichael and Bednarek, 2013; Müller and Jürgens, 2016). The phragmoplast expands by nucleation of new microtubules on pre-existing microtubules (Murata et al., 2013; Smertenko et al., 2018) and is partially dependent on the mitotic microtubule binding protein ENDOSPERM DEFECTIVE1 and the augmin complex to recruit gamma tubulin to phragmoplast microtubules (Lee et al., 2017; Nakaoka et al., 2012). Finally, the phragmoplast reaches the cell cortex and the cell plate and associated membranes fuse with the mother cell membranes at the cell plate fusion site previously specified by the PPB (van Oostende-Triplet et al., 2017).

TANGLED1 (TAN1, AT3G05330) was the first protein identified to localize to the plant division site throughout mitosis and cytokinesis (Walker et al., 2007). In maize, the *tan1* mutant has defects in division plane orientation caused by phragmoplast guidance defects (Cleary and Smith, 1998; Martinez et al., 2017). TAN1 bundles and crosslinks microtubules in vitro (Martinez et al., 2020). In vivo, TAN1 promotes microtubule pausing at the division site (Bellinger et al., 2021). TAN1, together with other division site localized proteins, is critical for the organization of an array of cell cortex localized microtubules that is independent from the phragmoplast. These cortical-telophase microtubules accumulate at the cell cortex during telophase and are subsequently incorporated into the phragmoplast to direct its movement towards the division site (Bellinger et al., 2021). Other important division site localized proteins were identified through their interaction with TAN1, such as the division site localized kinesin-12 proteins PHRAGMOPLAST ORIENTING KINESIN1 (POK1) and POK2 (Müller et al., 2006; Lipka et al., 2014). Similar to other kinesin-12 proteins, PHRAGMOPLAST ASSOCIATED KINESIN RELATED PROTEIN (PAKRP1) and PAKRPL1 (Lee et al., 2007; Pan et al., 2004), POK2 localizes to the phragmoplast midline during telophase and plays a unique role in phragmoplast expansion (Herrmann et al., 2018). Together, POK1 and POK2 are required to guide the phragmoplast to the division site (Herrmann et al., 2018; Müller et al., 2006). *pok1 pok2 Arabidopsis thaliana* (Arabidopsis) double mutants have stunted growth and misplaced cell walls as a result of phragmoplast guidance defects (Müller et al., 2006). The *pok1 pok2* double mutants also fail to maintain TAN1 at the division site after entry into metaphase (Lipka et al., 2014). This suggests that TAN1 maintenance at the division site after metaphase is dependent on POK1 and POK2.

In Arabidopsis, *tan1* mutants have very minor phenotypes (Walker et al., 2007). However, combination of *tan1* with *auxin-induced-in-root-cultures9* (*air9*), a mutant with no obvious defects (Buschmann et al. 2015), resulted in a synthetic phenotype consisting of reduced root growth, increased root cell file rotation and phragmoplast guidance defects (Mir et al. 2018). TAN1 and AIR9 are unrelated microtubule-binding proteins that both localize to the division site (Walker et al., 2007; Buschmann et al., 2006). Both TAN1 and AIR9 colocalize with the PPB. TAN1 remains at the division site throughout cell division, while AIR9 is lost from the division site upon PPB disassembly and then reappears at the division site during cytokinesis when the phragmoplast contacts the cortex. When full length *TAN1* fused to *YELLOW FLUORESCENT PROTEIN* (*TAN1-YFP*) and driven either by the constitutive viral cauliflower mosaic *CaMV35S* promoter (*p35S:TAN1-YFP*) or the native promoter with the fluorescent protein as either N- or C-terminal fusion (*pTAN1:TAN1-YFP* or *pTAN1:CFP-TAN1*) was transformed into the *tan1 air9* double mutant, the phenotype was rescued such that plants looked similar to and grew as well as wild-type plants (Mir et al., 2018; Mills and Rasmussen, 2022).

TAN1 is an intrinsically disordered protein with no well-defined domains. It was divided into five conserved regions based on alignments of amino acid similarity across plant species. Region I, which covers the first ~130 amino acids of TAN1, is the most highly conserved, and mediates TAN1 localization to the division site during telophase. This ~130 amino acid region also mediates interactions between TAN1 and POK1 in the yeast-two-hybrid system (Rasmussen et al., 2011). When *TAN1* missing the first ~130 amino acids was transformed into the *tan1 air9* double mutant, no rescue was observed (Mir et al., 2018). This suggests that the first ~130 amino acids of the TAN1 protein are critical for function in root growth and division plane positioning.

Here, we show that both AIR9 and TAN1 are required for POK1 to remain at the division site after PPB disassembly. We identified TAN1-POK1 interaction motifs within the first 132 amino acids using the yeast-two-hybrid system. Interestingly, the first 132 amino acids of TAN1 (TAN1_1-132_) are sufficient to rescue the *tan1 air9* double mutant, but not when a TAN1-POK1 interaction motif was disrupted. We found that when full-length TAN1 with the same mutated motif was used, substantial rescue was observed, except defects in phragmoplast guidance and loss of POK1 and TAN1 at the division site during metaphase and telophase. Together, this suggests that interactions between POK1 and AIR9, and TAN1 and POK1, as well as other yet unknown proteins, are important for division plane orientation and plant growth.

## Results

### Either TAN1 or AIR9 is sufficient to recruit and maintain POK1 at the division site

To understand how known division-site localized proteins interact at the division site, we examined POK1 fused to YELLOW FLUORESCENT PROTEIN (YFP-POK1 (Lipka et al., 2014)) localization in wild type, *tan1 air9* double mutants and single mutants expressing the microtubule marker *UBQ10:mScarlet-MAP4* (Pan et al., 2020). Our hypothesis was that POK1 localization would not be contingent on TAN1 or AIR9, and would therefore be unaltered in the *tan1 air9* double mutant. In contrast to our hypothesis, YFP-POK1 was lost from the division site during metaphase and telophase and also accumulated less frequently during preprophase and prophase. In wild-type cells, YFP-POK1 colocalized with PPBs in 71% of preprophase/prophase cells (n = 50/70 cells, 15 plants, Figure 1A) consistent with previous observations (Schaefer et al., 2017). In the *tan1 air9* double mutant, YFP-POK1 colocalized with 50% of PPBs during preprophase/prophase, which was not significantly different than wild-type (n = 27/54 cells, 15 plants, Figure 1D; Table 1; Fisher’s exact test, P-Value = 0.0165, ns with Bonferroni correction). In wild-type cells, YFP-POK1 remained at the division site in all observed metaphase (n = 13/13 cells, Figure 1B) and telophase cells (n = 31/31 cells, Figure 1C), similar to previous studies (Lipka et al., 2014). In rare instances, YFP-POK1 also accumulated in the phragmoplast midline in wild-type cells (13%, n = 4/31, 11 plants, Table 1). In contrast, in *tan1 air9* mutants, YFP-POK1 was lost from the division site in metaphase (n = 0/21 cells, Figure 1E; Table 1) and telophase (n = 0/44, Figure 1F). Interestingly, in *tan1 air9* double mutants, although YFP-POK1 did not accumulate at the division site, it accumulated at the phragmoplast midline in 77% of cells (n = 34/44), significantly more frequent midline accumulation than the 13% observed in wild-type plants (n = 4/31 cells, Table 1; Fisher’s exact test, P-value < 0.00001). Together, this shows that POK1 is not maintained at the division site after PPB disassembly and that instead it accumulates in the phragmoplast midline. We hypothesize that mislocalized phragmoplast midline accumulation of YFP-POK1 in *tan1 air9* mutants occurs because YFP-POK1 is not maintained at the division site.

**Figure 1:**
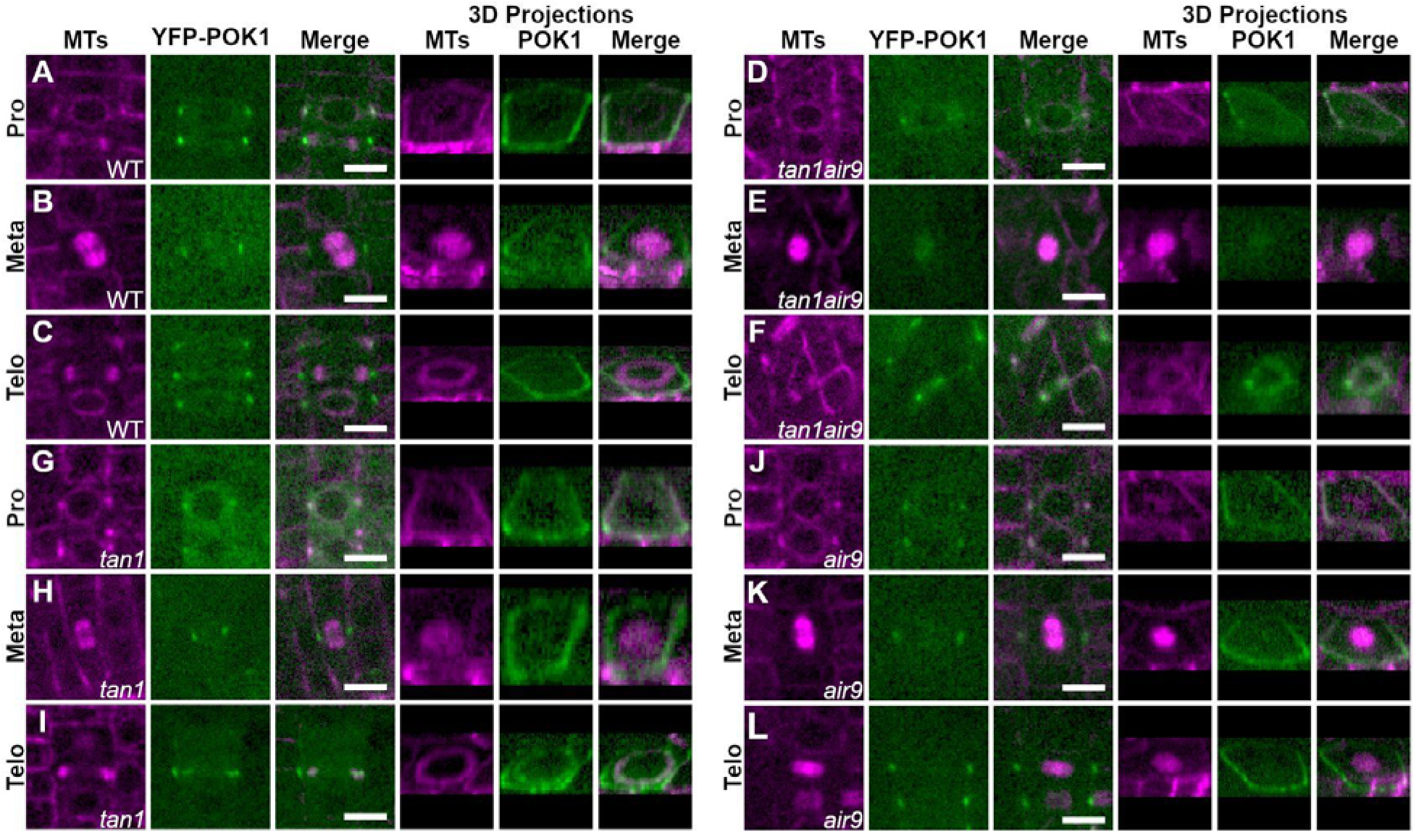
TAN1 and AIR9 together promote POK1 maintenance at the division site. YFP-POK1 localization in Col-0 wild-type plants, *tan1* single mutants, *air9* single mutants, and *tan1 air9* double mutants expressing *UBQ10:mScarlet-MAP4* to mark microtubules and *pPOK1:YFP-POK1*. Scale bars = 10 μm. A-C) YFP-POK1 localization in Col-0 wild-type plants, N = 15 plants. A) YFP-POK1 localization during preprophase/prophase. In 71% of cells (50/70) YFP-POK1 colocalized with the PPB. B) YFP-POK1 was observed to be maintained at the division site in metaphase and anaphase cells in Col-0 plants (n = 13/13 metaphase cells, n = 3/3 anaphase cells). C) YFP-POK1 remains clearly visible at the division site in Col-0 telophase cells (n = 31/31 cells). D-F) YFP-POK1 localization in *tan1 air9* double mutant plants, N = 19 plants. D) YFP-POK1 localization during preprophase/prophase. In 50% of cells (27/54) YFP-POK1 colocalized with the PPB. E) YFP-POK1 was observed to be lost from the division site upon entry into metaphase (n = 0/21 cells) and was absent in anaphase cells (n = 0/4 cells). F) In *tan1 air9* telophase cells YFP-POK1 was absent from the division site and accumulated in the phragmoplast midline (n = 34/44 cells). G-I) YFP-POK1 localization in *tan1* single mutants, N = 17 plants. G) YFP-POK1 localization during preprophase/prophase. In 64% of cells (54/85) YFP-POK1 colocalized with the PPB. H) YFP-POK1 was observed to be maintained at the division site in metaphase and anaphase cells in *tan1* plants (n = 17/17 metaphase cells, n = 6/6 anaphase cells). I) YFP-POK1 remains clearly visible at the division site in *tan1* telophase cells (n = 27/27 cells). J-L) YFP-POK1 localization in *air9* single mutant plants (N = 15 plants). J) YFP-POK1 localization during preprophase/prophase. In 64% of cells (46/72) YFP-POK1 colocalized with the PPB. K) YFP-POK1 was maintained at the division site in metaphase and anaphase cells in *air9* plants (n = 24/24 metaphase cells, n = 4/4 anaphase cells). L) YFP-POK1 remains clearly visible at the division site in *air9* telophase cells (n = 40/40 cells).

**Table 1.**
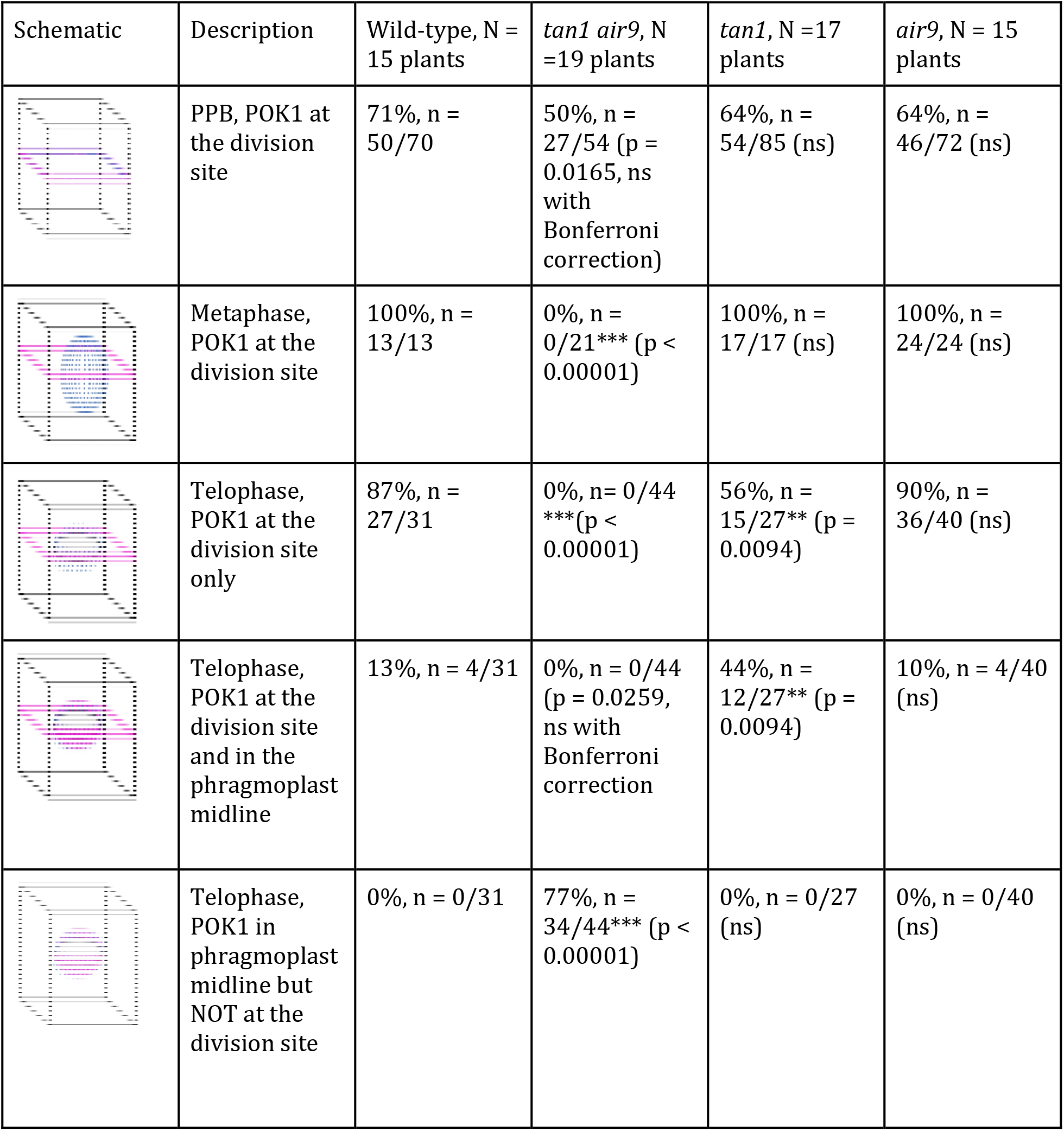
YFP-POK1 division site and phragmoplast midline accumulation in wild-type, *tan1, air9*, and *tan1 air9* double mutant plants. Statistically significant differences were determined using Fisher’s exact test with Bonferroni correction - for 4 sample types, P < 0.0125 is significant. P-values are in parentheses. Stars indicate significant differences; ns indicates not significant. Magenta represents YFP-POK1, blue represents microtubules, and light gray represents the cell plate in schematics.

| Schematic | Description | Wild-type, N = 15 plants | *tan1 air9*, N = 19 plants | *tan1*, N = 17 plants | *air9*, N = 15 plants |
| --- | --- | --- | --- | --- | --- |
|  | PPB, POK1 at the division site | 71%, n = 50/70 | 50%, n = 27/54 (p = 0.0165, ns with Bonferroni correction) | 64%, n = 54/85 (ns) | 64%, n = 46/72 (ns) |
|  | Metaphase, POK1 at the division site | 100%, n = 13/13 | 0%, n = 0/21*** (p < 0.00001) | 100%, n = 17/17 (ns) | 100%, n = 24/24 (ns) |
|  | Telophase, POK1 at the division site only | 87%, n = 27/31 | 0%, n = 0/44 *** (p < 0.00001) | 56%, n = 15/27** (p = 0.0094) | 90%, n = 36/40 (ns) |
|  | Telophase, POK1 at the division site and in the phragmoplast midline | 13%, n = 4/31 | 0%, n = 0/44 (p = 0.0259, ns with Bonferroni correction) | 44%, n = 12/27** (p = 0.0094) | 10%, n = 4/40 (ns) |
|  | Telophase, POK1 in phragmoplast midline but NOT at the division site | 0%, n = 0/31 | 77%, n = 34/44*** (p < 0.00001) | 0%, n = 0/27 (ns) | 0%, n = 0/40 (ns) |

Next, we examined YFP-POK1 localization in *tan1* and *air9* single mutants. YFP-POK1 localized to the division site during all mitotic stages, but aberrantly accumulated in the phragmoplast midline in *tan1* single mutants. Similar to wild-type plants, YFP-POK1 colocalized with PPBs during preprophase or prophase in *tan1* mutants (Figure 1G) and *air9* mutants (Figure 1H), and remained at the division site during metaphase and telophase (Figure 1G-L, Table 1). In *tan1* single mutants, YFP-POK1 localized both to the division site and the phragmoplast midline in 44% of telophase cells (Figure 1I, n = 12/27), which is significantly more midline accumulation compared to wild-type plants (13%, n = 4/31, 10 plants, Fisher’s exact test, P-value = 0.0094) or *air9* single mutants (Figure 1L, 10%, n = 4/40). Aberrant phragmoplast midline accumulation of YFP-POK1 in the *tan1* single mutants suggested that POK1-TAN1 interaction might be required to maintain POK1 at the division site. This prompted us to examine their interaction more closely.

### Amino acids 1-132 of TAN1 Rescue the *tan1 air9* Double Mutant

POK1 interacts with both full-length TAN1 and the first 132 amino acids of TAN1 using the yeast-two-hybrid system (Rasmussen et al., 2011). In addition, TAN1 missing the first 126 amino acids failed to rescue the *tan1 air9* double mutant, suggesting that this part of the protein is critical for TAN1 function (Mir et al., 2018). To test the function of this region of the protein in Arabidopsis, the *TAN1* coding sequence for the first 132 amino acids was fused to YFP (TAN1_1-132_-YFP) driven by the cauliflower mosaic *p35S* promoter and was then transformed into the *tan1 air9* double mutant. We used *p35S:TAN1-YFP* in the *tan1 air9* double mutant as our benchmark for rescue, as its ability to rescue the *tan1 air9* double mutant was demonstrated previously (Mir et al., 2018). The progeny of several independent *p35S:TAN1_1-132_-YFP* lines rescued the *tan1 air9* double mutant, as described in more detail below. Overall root patterning of *tan1 air9* double mutants expressing either *p35S:TAN1_1-132_-YFP* or full-length *p35S:TAN1-YFP* was restored, while untransformed *tan1 air9* double mutant roots had misoriented divisions (Figure 2A, Supplementary Figure 1). Cell file rotation, which skews left and has large variance in the *tan1 air9* double mutant (Figure 2B & 2C), was significantly rescued in both *p35S:TAN1_1-132_-YFP* and *p35S:TAN1-YFP tan1 air9* lines (n = 37 and 41 plants respectively), compared to the untransformed *tan1 air9* control (Levene’s test used due to non-normal distribution, P-value < 0.0001). Root length at 8 days after stratification was also restored (Figure 2D). Interestingly, although TAN1_1-132_-YFP rarely co-localizes with PPBs in wild-type plants (Rasmussen et al., 2011) or in *tan1 air9* double mutants (10%, n = 9/89 cells, Figure 3A), PPB angles of *p35S:TAN1_1-132_-YFP* and *p35S:TAN1-YFP tan1 air9* plants had significantly less variance compared to the untransformed control (Figure 2E). Phragmoplast positioning defects of the *tan1 air9* double mutant were also significantly rescued by *p35S:TAN1_1-132_-YFP*. Altogether, *p35S:TAN1_1-132_-YFP* rescued the phenotypes of the double mutant similar to full-length *p35S:TAN1-YFP*. This indicates that most functions that affect phenotypes assessed here are encoded by the first section of the *TAN1* gene.

**Figure 2.**
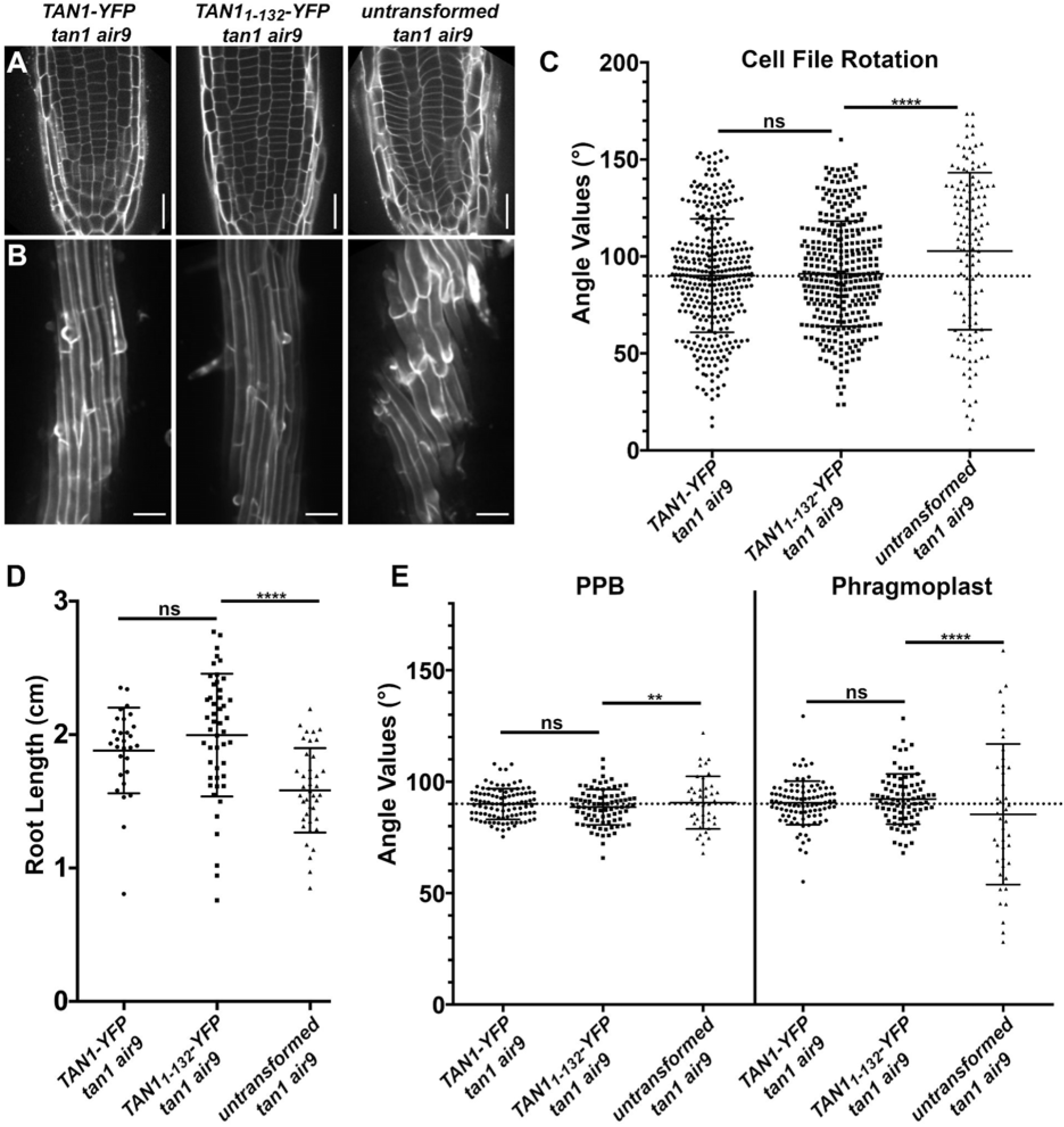
*p35S:TAN1_1-132_-YFP* rescues Arabidopsis *tan1 air9* double mutant phenotypes. A) Cell walls stained with propidium iodide (PI) of *tan1 air9* double mutant root tips expressing *p35S:TAN1-YFP* (left), *p35S:TAN1_1-132_-YFP* (middle), and untransformed *tan1 air9* double mutants (right). Bars = 25 μm. B) Maximum projections of 10 1-μm Z-stacks of PI-stained differentiation zone root cell walls. Scale bars = 50 μm. C) Cell file rotation angles of *tan1 air9* double mutants expressing *p35S:TAN1-YFP* (left), *p35S:TAN1_1-132_-YFP* (middle) and untransformed plants (right), n > 13 plants for each genotype. Each dot represents an angle measured from the left side of the long axis of the root to the transverse cell wall. Angle variances were compared with Levene’s test due to non-normal distribution. D) Root length measurements from 8 days after stratification of *tan1 air9* double mutants expressing *p35S:TAN1-YFP* (left), *p35S:TAN1_1-132_-YFP* (middle) and untransformed plants (right), n > 28 plants for each genotype, compared by two-tailed t-test with Welch’s correction. E) PPB and phragmoplast angle measurements in *tan1 air9* double mutant cells expressing *p35S:TAN1-YFP* (left), *p35S:TAN1_1-132_-YFP* (middle) and untransformed plants (right), n > 20 plants for each genotype. Angle variations compared with F-test. ns indicates not significant, ** P-value <0.01, **** P-value <0.0001.

**Figure 3:**
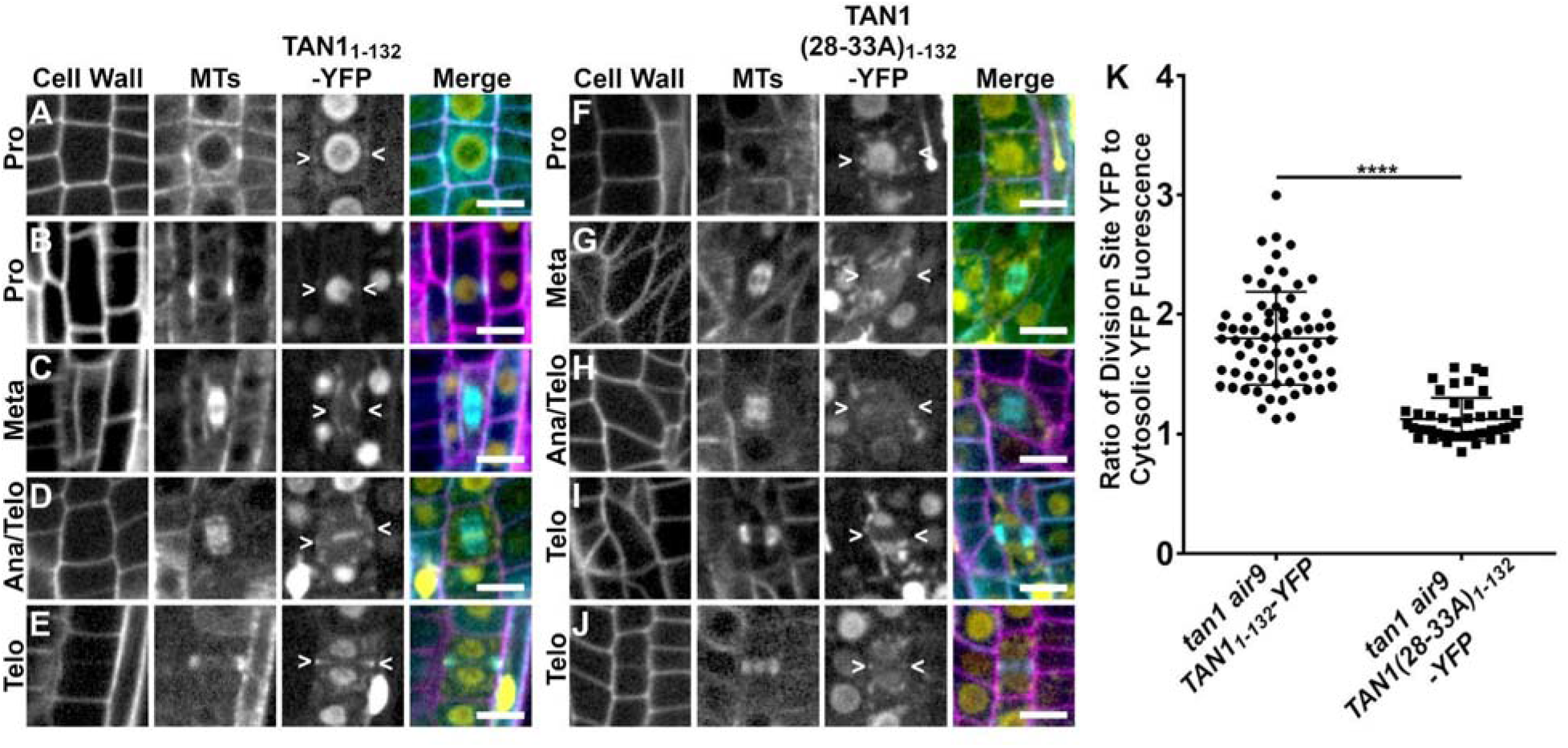
Division site localization during telophase is common for TAN1_1-132_-YFP but rare for TAN1(28-33A)_1-132_-YFP in *tan1 air9* double mutants. A-E) Propidium iodide stained *tan1 air9* plants expressing *p35S:TAN1_1-132_-YFP* during mitosis (n = 29 plants). The division site is indicated by arrowheads in the YFP panels. Scale bars = 10 μm. A) Rare prophase division site accumulation of TAN1_1-132_-YFP, (10%, n = 9/89 cells), (B) common prophase TAN1_1-132_-YFP nuclear accumulation without division site localization (90%, n = 80/89 cells), (C) no specific TAN1_1-132_-YFP division site accumulation in metaphase (100%, n = 28/28 cells), (D) faint TAN1_1-132_-YFP division site accumulation accompanied by midline accumulation in late anaphase/early telophase (80%, n = 16/20 cells) and E) TAN1_1-132_-YFP division site accumulation during telophase (100%, n = 58/58 cells). F-H) *tan1 air9* plants expressing *p35S:TAN1(28-33A)_1-132_-YFP* during mitosis (n = 13 plants). The division site is indicated by arrowheads in the YFP panels. F) No specific TAN1(28-33A)_1-132_-YFP prophase division site accumulation during prophase (100%, n = 20/20 cells), (G) no specific TAN1(28-33A)_1-132_-YFP division site accumulation during metaphase (100%, n = 12/12 cells), (H) no TAN1(28-33A)_1-132_-YFP division site or midline accumulation in late anaphase/early telophase (100%, n = 8/8 cells), (I) no specific TAN1(28-33A)_1-132_-YFP division site accumulation during telophase (68%, n = 15/22 cells) and (J) faint TAN1(28-33A)_1-132_-YFP division site accumulation during telophase (32%, n = 7/22 cells). K) Ratio of TAN1_1-132_-YFP (left) or TAN1(28-33A)_1-132_-YFP (right) fluorescence at the division site to cytosolic fluorescence from *tan1 air9* plants expressing *p35S:TAN1_1-132_-YFP* or *p35S:TAN1(28-33A)_1-132_-YFP* during telophase, n >23 plants for each genotype. Asterisks indicate a significant difference as determined by Mann-Whitney U test, P-value <0.0001.

### Disrupting TAN1-POK1 interaction alters TAN1 and POK1 localization to the division site and reduces *tan1 air9* rescue

To further understand how TAN1 functions, we disrupted its ability to interact with the kinesin POK1 using alanine scanning mutagenesis. Alanine scanning mutagenesis was used to replace six amino acids with six alanines across the first ~120 amino acids of TAN1_1-132_ (described in materials and methods). After testing their interaction with POK1 using the yeast-two-hybrid system, we identified seven constructs that lost interaction with POK1 (Supplementary Figure 2). Reasoning that highly conserved amino acids would be more likely to play critical roles in TAN1-POK1 interaction, we selected TAN1_1-132_ with alanine substitutions replacing the highly conserved amino acids 28-33 (INKVDK) with six alanines (TAN1(28-33A)_1-132_) for analysis in Arabidopsis. Our hypothesis was that the mutated form of TAN1_1-132_ (TAN1(28-33A)_1-132_) would not rescue the *tan1 air9* mutant due to lack of POK1 and TAN1 interaction. *TAN1(28-33A)_1-132_* was cloned into a plant transformation vector to generate *p35S:TAN1(28-33A)_1-132_-YFP* and transformed into the *tan1 air9* double mutant. The *p35S:TAN1(28-33A)_1-132_-YFP* construct partially rescued the *tan1 air9* double mutant (Figure 4). *p35S:TAN1(28-33A)_1-132_-YFP* in the *tan1 air9* double mutant did not rescue cell file rotation defects (Figure 4B, D) or phragmoplast angle defects (Figure 4F). However, overall plant growth (Figure 4C) and root length (Figure 4E) showed intermediate rescue compared to unaltered *p35S:TAN1_1-132_-YFP* in the *tan1 air9* double mutant. PPB angles in *tan1 air9* double mutants expressing either *p35S:TAN1(28-33A)_1-132_-YFP* or *p35S:TAN1_1-132_-YFP* were similar, suggesting that TAN1-POK1 interaction may not be required for PPB placement (Figure 4F). These results suggest that the first 132 amino acids of TAN1 perform several vital functions, some of which are contingent or partially contingent on a likely interaction with POK1 in Arabidopsis.

**Figure 4:**
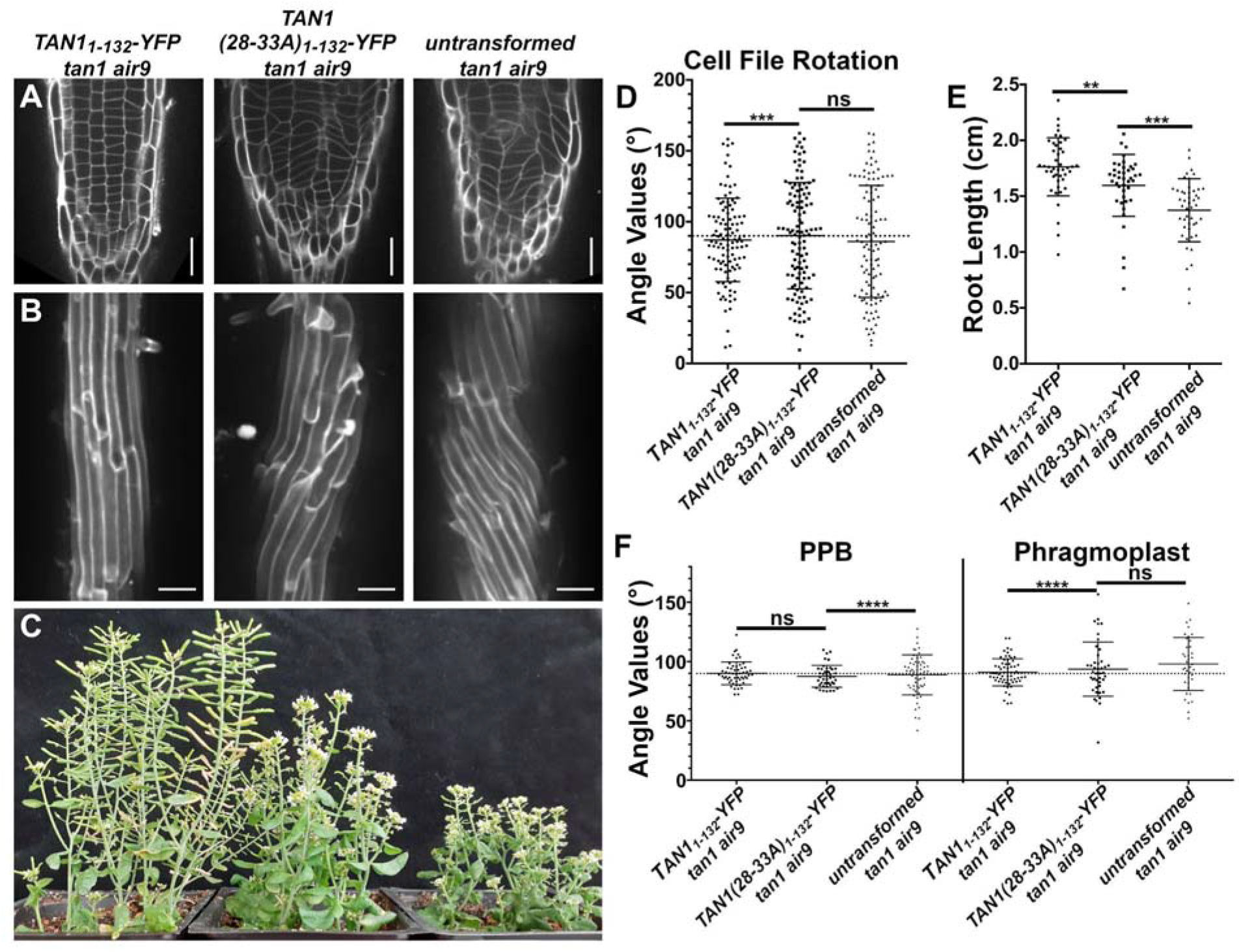
*p35S:TAN1(28-33A)_1-132_-YFP* partially rescues *tan1 air9* double mutant phenotypes. A) Cell walls of Arabidopsis *tan1 air9* double mutant root tips stained with propidium iodide (PI) of plants expressing *p35S:TAN1_1-132_-YFP* (left), *p35S:TAN1(28-33A)_1-132_-YFP* (middle), and untransformed *tan1 air9* double mutants (right). Scale bars = 25 μm. B) Maximum projections of 10 1-μm Z-stacks of PI-stained differentiation zone root cell walls. Scale bars = 50 μm. C) 58-day old *tan1 air9* double mutants expressing *p35S:TAN1_1-132_-YFP* (left), *p35S:TAN1(28-33A)_1-132_-YFP* (middle), and untransformed *tan1 air9* double mutants (right). D) Cell file rotation angles of *tan1 air9* double mutants expressing *p35S:TAN1_1-132_-YFP* (left), *p35S:TAN1(28-33A)_1-132_-YFP* (middle), and untransformed *tan1 air9* double mutants (right) n > 27 plants for each genotype. Variances were compared with Levene’s test. E) Root length measurements from 8 days after stratification of *tan1 air9* double mutants expressing *p35S:TAN1_1-132_-YFP* (left), *p35S:TAN1(28-33A)_1-132_-YFP* (middle), and untransformed *tan1 air9* double mutants (right), n > 40 plants for each genotype, two-tailed t-test with Welch’s correction. F) PPB and phragmoplast angle measurements in dividing root cells of *tan1 air9* double mutants expressing *p35S:TAN1_1-132_-YFP* (left), *p35S:TAN1(28-33A)_1-132_-YFP* (middle), and untransformed plants (right), n > 17 plants for each genotype. Angle variance compared with F-test. ns indicates not significant, ** P-value <0.01, *** P-value <0.001, **** P-value <0.0001.

To understand how this mutation within TAN1*_1-132_* affected localization, we analyzed TAN1(28-33A)_1-132_-YFP in the *tan1 air9* double mutant. Localization of TAN1(28-33A)_1-132_-YFP to the division site in *tan1 air9* double mutants was significantly reduced compared to unaltered TAN1_1-132_-YFP, which localized to the division site during telophase 100% of the time (n = 58/58 cells, 29 plants, Figures 3E, (Rasmussen et al., 2011)). TAN1(28-33A)_1-132_-YFP showed no obvious division site localization 68% of the time (n = 15/22 cells, Figure 3I) or faint division site accumulation in 32% of telophase cells (n = 7/22 cells, Figure 3J). When the fluorescence intensity of TAN1(28-33A)_1-132_-YFP at the division site during telophase was compared to the cytosolic fluorescence intensity in the same cell, the median ratio was ~1.1 indicating little preferential accumulation of TAN1(28-33A)_1-132_-YFP at the division site (Figure 3K). In contrast, the median ratio of unaltered TAN1_1-132_-YFP at the division site was ~1.8 compared to cytosolic fluorescence, indicating its preferential accumulation at the division site. This suggests that TAN1 requires the motif in amino acids 28-33 to localize properly to the division site during telophase. Our hypothesis is that this reduced localization is due to disruptions in TAN1_1-132_-POK1 interaction.

Next, we generated a construct that introduced alanines at amino acids 28-33 in full-length YFP-TAN1 constructs (*p35S:YFP-TAN1(28-33A)*) to assess whether *p35S:YFP-TAN1(28-33A)* would rescue the *tan1 air9* double mutant. In contrast to the modest partial rescue provided by *p35S:TAN1(28-33A)_1-132_-YFP*, full-length *p35S:YFP-TAN1(28-33A)* significantly rescued the defects in the *tan1 air9* double mutant, as described below. First, we assessed whether full-length TAN1(28-33A) interacted with POK1 via the yeast-two-hybrid system, and it did not (Supplementary Figure 3). Next, we analyzed rescue in Arabidopsis expressing *p35S:YFP-TAN1(28-33A)*. Most defects except phragmoplast angle variance (Figure 5, Supplementary Figure 4) were fully rescued in the *p35S:YFP-TAN1(28-33A) tan1 air9* lines, including cell file rotation (Figure 5C), root length (Figure 5D) and PPB angles (Figure 5E). Similar to TAN1-YFP, YFP-TAN1(28-33A) localized to the division site in preprophase, prophase and telophase (Supplementary Figure 5).

**Figure 5:**
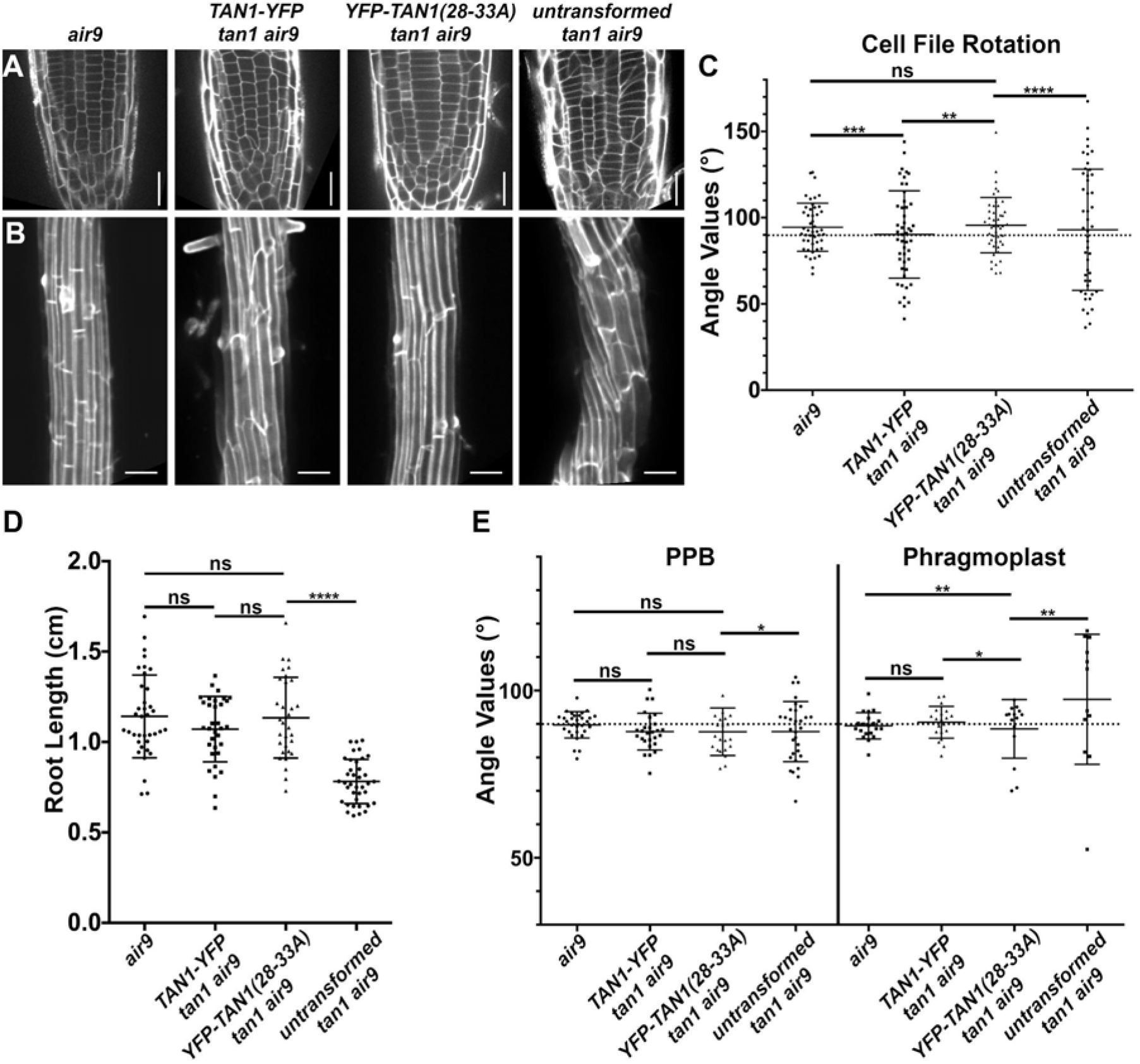
Full length *TAN1* with alanine substitutions replacing amino acids 28 to 33 (*p35S:YFP-TAN1(28-33A)*) mostly rescues the t*an1 air9* double mutant. A) Propidium iodide stained root tips of an *air9* single mutant (left) and *tan1 air9* double mutants expressing *p35S:TAN1-YFP* (center left) or *p35S:YFP-TAN1(28-33A)* (center right), and an untransformed *tan1 air9* plant (right). Scale bars = 25 μm. B) Maximum projections of 10 1-μm Z-stacks of PI-stained cell walls in the root differentiation zone. Scale bars = 50 μm. C) Cell file rotation angles of *air9* single mutants (left), *tan1 air9* double mutant plants expressing *p35S:TAN1-YFP* (center left) or *p35S:YFP-TAN1(28-33A)* (center right), and untransformed *tan1 air9* plants (right), n > 9 plants for each genotype. Variances were compared with Levene’s test. D) Root length measurements from 8 days after stratification of *air9* single mutants (left) and *tan1 air9* double mutants expressing *p35S:TAN1-YFP* (center left) or *p35S:YFP-TAN1(28-33A)* (center right), and untransformed *tan1 air9* plants (right), n > 30 plants of each genotype, compared by two-tailed t-test with Welch’s correction. E) PPB and phragmoplast angle measurements in dividing root cells of *air9* single mutants (left) and *tan1 air9* double mutant plants expressing *p35S:TAN1-YFP* (center left) or *p35S:YFP-TAN1(28-33A)* (center right), and untransformed *tan1 air9* plants (right), PPB measurements n > 15 plants for each genotype; phragmoplast measurements n > 8 plants for each genotype. Angle variance compared with F-test. ns indicates not significant, * P-value <0.05, ** P-value <0.01, *** P-value <0.001, **** P-value <0.0001.

To determine if full-length YFP-TAN1(28-33A) had reduced accumulation at the division site during telophase similar to TAN1(28-33A)_1-132_-YFP, fluorescence intensity levels were measured. During prophase, YFP-TAN1(28-33A) fluorescence intensity at the division site compared to the cytosol was comparable to TAN1-YFP fluorescence intensity ratios. In contrast, YFP-TAN1(28-33A) fluorescence intensity ratios during telophase were reduced to ~1.6 compared with unaltered TAN1-YFP (~2.1) indicating that YFP-TAN1(28-33A) accumulated less at the division site during telophase (Supplementary Figure 5). Together, these data suggest that TAN1 is recruited to the division site during prophase without interaction with POK1. Defects in phragmoplast positioning may be due specifically to the disruption of TAN1-POK1 interaction, or due to the lower accumulation of TAN1 at the division site that would normally be mediated by POK1 during telophase.

To better understand how these alanine substitutions affect both POK1 and TAN1 localization we examined *tan1 air9* double mutants expressing a microtubule marker (*UBQ10:mScarlet-MAP4* (Pan et al., 2020)), *pTAN1:CFP-TAN1(28-33A)* and *pPOK1:YFP-POK1* (Lipka et al. 2014). Both CFP-TAN1(28-33A) and YFP-POK1 had reduced accumulation at the division site in the *tan1 air9* double mutant. CFP-TAN1(28-33A) and YFP-POK1 colocalized with the PPB in 41% of cells (n = 32/79) which is significantly less frequent when compared to 72% of cells (n=58/82 cells, 20 plants) with PPBs in *tan1 air9* mutants containing unaltered CFP-TAN1 and YFP-POK1 (Figure 6A, Table 2; Fisher’s exact test, P-value = 0.0001). Unaltered CFP-TAN1 fully rescued the *tan1 air9* double mutant (Mills and Rasmussen, 2022), and serves here as a control. Unaltered CFP-TAN1 and YFP-POK1 localized and were maintained at the division site similar to wild type in metaphase (Figure 6B, n=13/13), while CFP-TAN1(28-33A) and YFP-POK1 in *tan1 air9* mutants were sometimes absent from the division site in metaphase with only 58% of metaphase cells maintaining both proteins at the division site (n = 11/19 cells, Figure 6F, Table 2). During early telophase, unaltered CFP-TAN1 and YFP-POK1 were always at the division site (n = 14/14, Figure 6C), but CFP-TAN1(28-33A) and YFP-POK1 were maintained at the division site in only 65% of early telophase cells (n = 20/31 cells, Figure 6G, Table 2). Interestingly, YFP-POK1 accumulated in the phragmoplast midline in 26% of early telophase cells (n = 8/31 cells, Table 2) but was not observed in the phragmoplast midline in early telophase cells of plants expressing unaltered CFP-TAN1 (n = 0/14 cells, Table 2). During late telophase, when the phragmoplast has contacted the cell cortex in at least one location, CFP-TAN1 and POK1 always localized to the division site (100%, n =63/63 cells, Figure 6D). Interestingly, although not observed in earlier stages, YFP-POK1 and CFP-TAN1(28-33A) recruitment to the division site increased to 90% of late telophase cells (n = 53/59 cells, Figure 6H). In the remainder of cells, neither CFP-TAN1(28-33) nor YFP-POK1 localized to the division site (3%, n = 2/59), or only CFP-TAN1(28-33) accumulated at the division site (7%, n = 4/59 cells). Together, these data suggest that TAN1-POK1 interactions play a critical role in stabilizing them together at the division site. Additionally, it suggests that other, yet unidentified proteins may recruit both TAN1 and POK1 to the division site, particularly during late telophase, in the absence of both AIR9 and TAN1-POK1 interaction.

**Figure 6:**
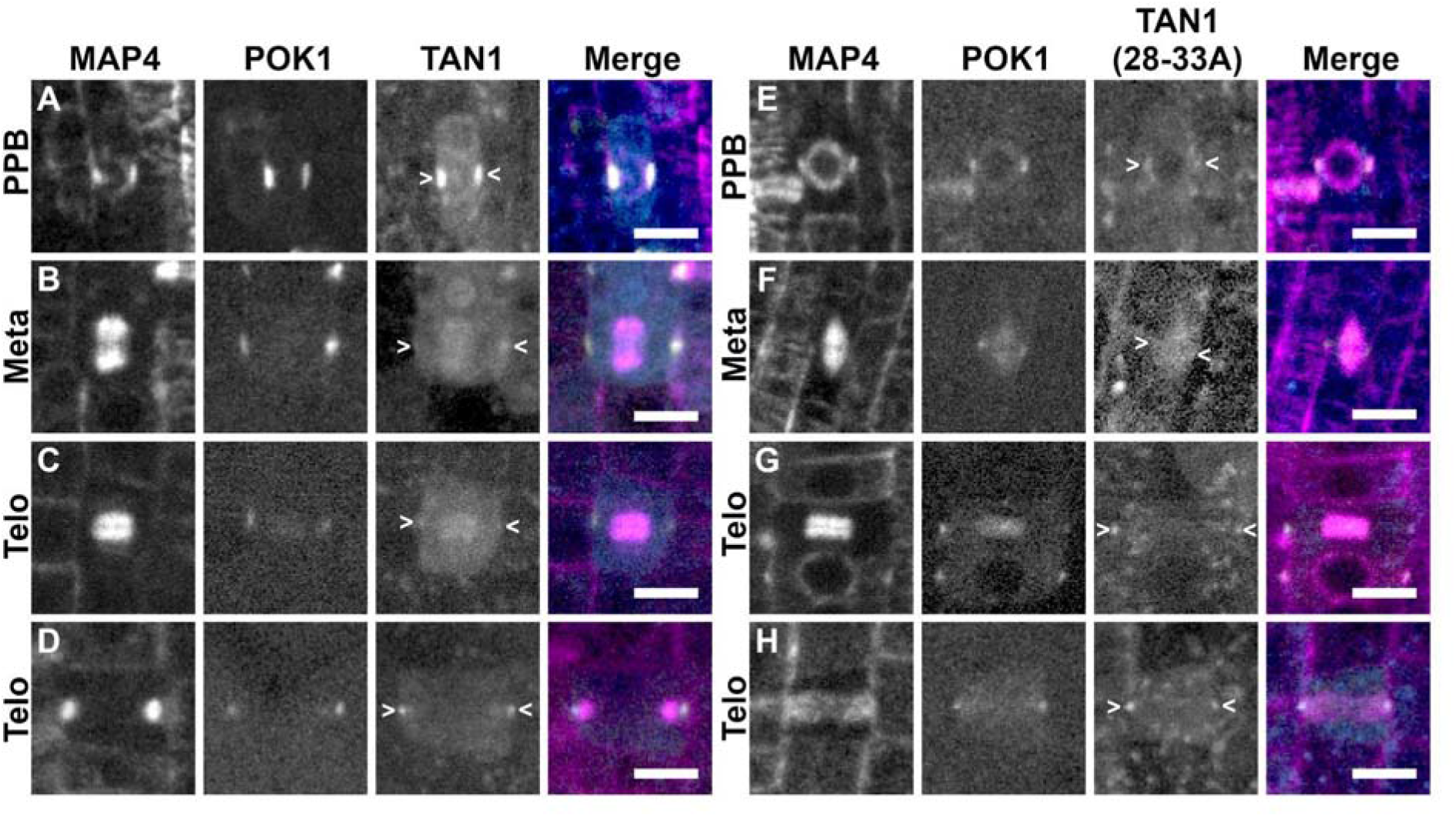
CFP-TAN1(28-33A) and YFP-POK1 exhibit impaired recruitment to the division site in the *tan1 air9* double mutant. YFP-POK1 localization in *tan1 air9* double mutants expressing *UBQ10:mScarlet-MAP4* and either (A-D) *pTAN1:CFP-TAN1* (n = 20 plants) or (E-I) *pTAN1:CFP-TAN1(28-33A)* (n = 22 plants). Maximum projections of 3 1-μm Z-stacks. Scale bars = 10 μm. Some bleed through from the mScarlet channel can be seen in the YFP-POK1 panels. A) YFP-POK1 and CFP-TAN1 colocalized with the PPB in 72% of cells (n = 59/82 cells). B) YFP-POK1 and CFP-TAN1 were maintained at the division site in metaphase 13/13 and anaphase 4/4 cells. C) YFP-POK1 and CFP-TAN1 were maintained at the division site in all early telophase cells (n = 14/14 cells) and late telophase (n = 63/63 cells). E) YFP-POK1 and CFP-TAN1(28-33A) colocalized with the PPB in 41% of cells (n = 32/79 cells). F) YFP-POK1 and CFP-TAN1(28-33A) were maintained at the division site in 58% of metaphase cells (n = 11/19 cells). CFP-TAN1(28-33A) was faint at the division site. G) Both YFP-POK1 and CFP-TAN1(28-33A) were observed at the division site in 65% of early telophase cells (n = 20/31 cells). H) YFP-POK1 and CFP-TAN1(28-33A) were recruited to the division site in 90% of late telophase cells (n = 53/59 cells). Some late telophase cells were observed to have CFP-TAN1(28-33A) but not YFP-POK1 at the division site (7%, n = 4/59 cells) or neither CFP-TAN1(28-33A) or YFP-POK1 at the division site (3% n = 2/59 cells).

**Table 2.**
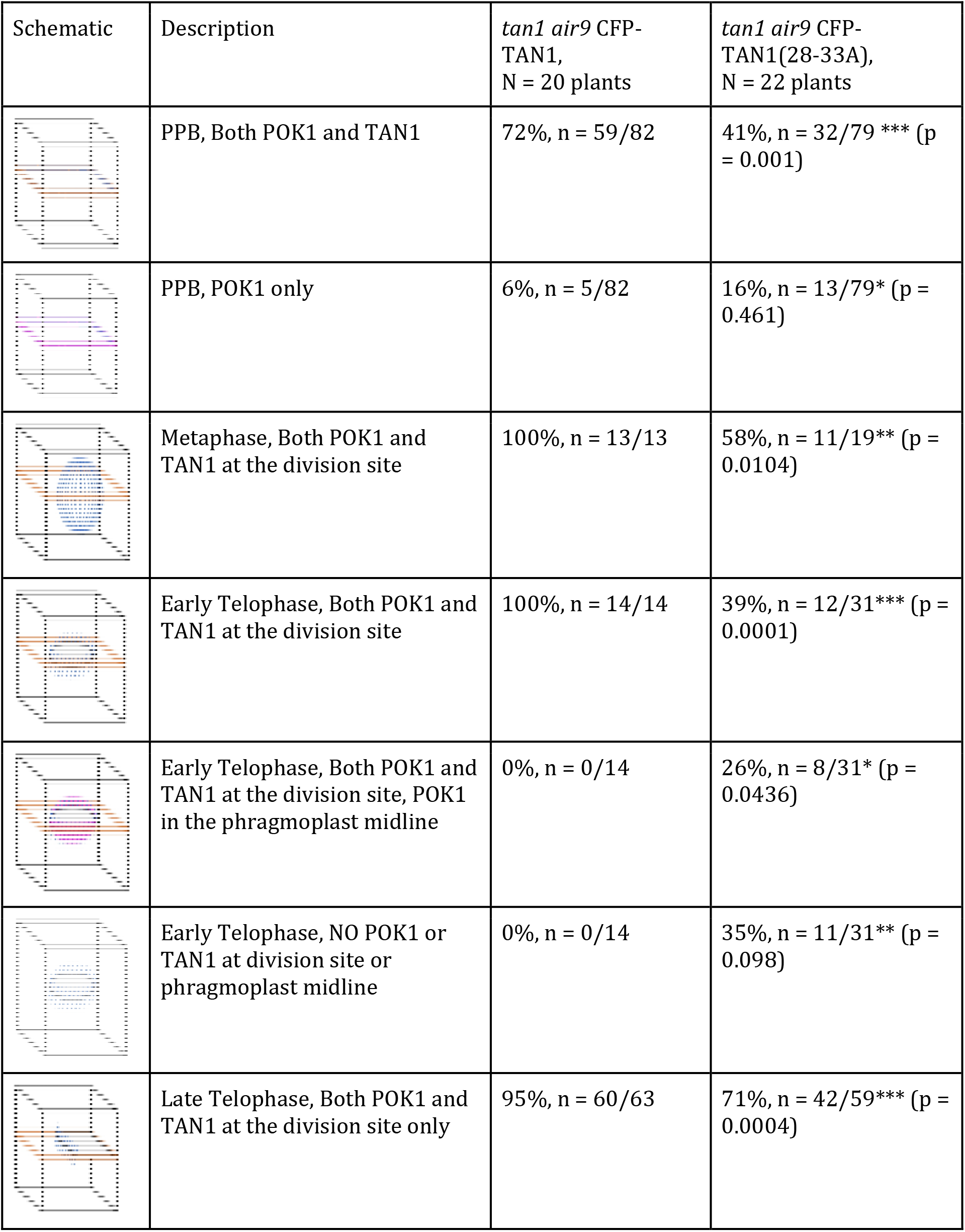
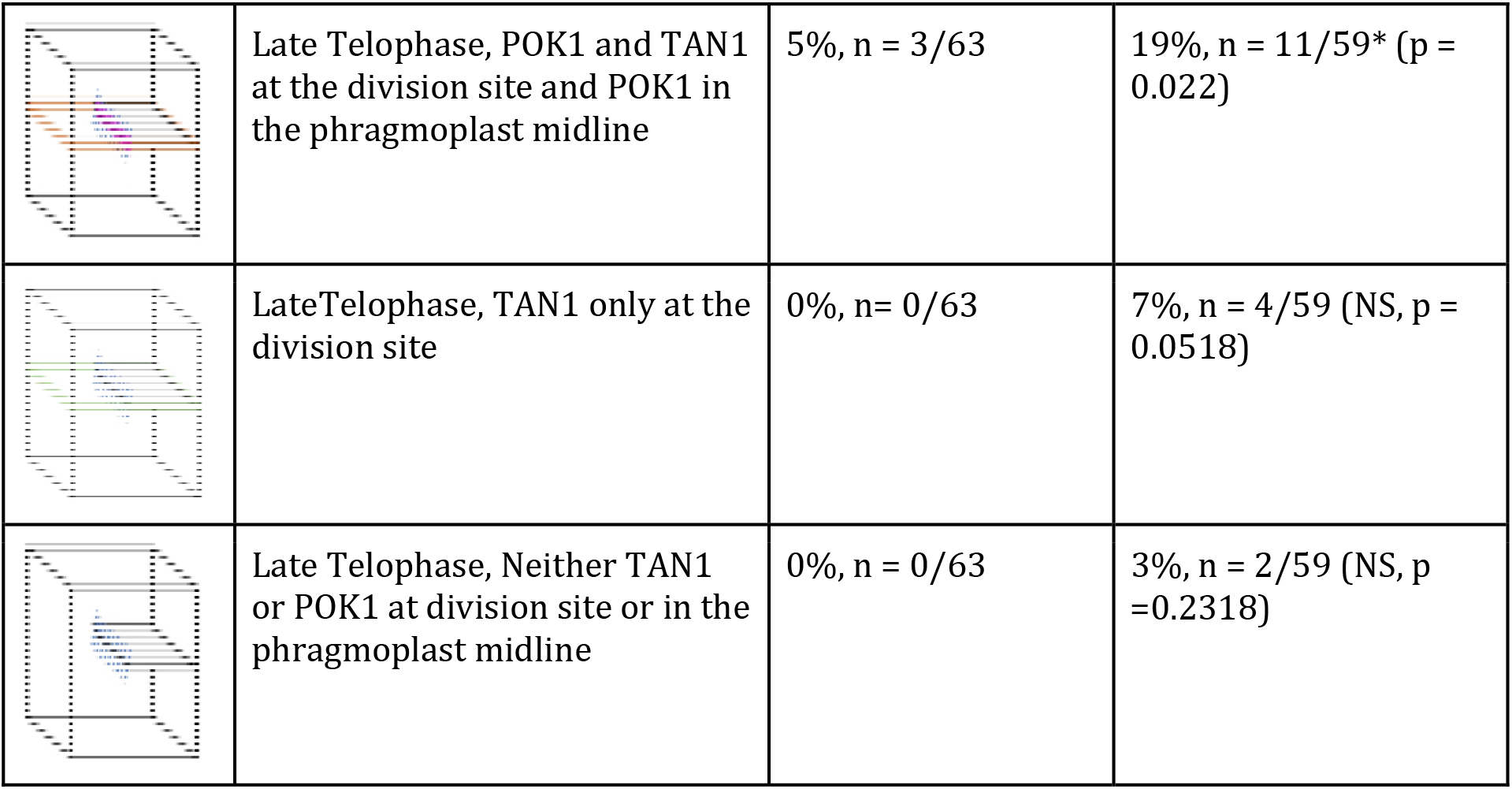
POK1 and TAN1 or TAN1(28-33) localization to the division site in *tan1 air9* double mutants. Statistically significant differences were determined using Fisher’s exact test. NS indicates not significant. Brown represents dual localization of YFP-POK1 and either CFP-TAN1 or CFP-TAN1(28-33A), magenta is YFP-POK1 alone, green is CFP-TAN1(28-33A) alone, blue lines are microtubules, and light gray represents the cell plate in schematics.

| Schematic | Description | *tan1 air9* CFP-TAN1,<br>N = 20 plants | *tan1 air9* CFP-TAN1(28-33A),<br>N = 22 plants |
| --- | --- | --- | --- |
|  | PPB, Both POK1 and TAN1 | 72%, n = 59/82 | 41%, n = 32/79 *** (p = 0.001) |
|  | PPB, POK1 only | 6%, n = 5/82 | 16%, n = 13/79* (p = 0.461) |
|  | Metaphase, Both POK1 and TAN1 at the division site | 100%, n = 13/13 | 58%, n = 11/19** (p = 0.0104) |
|  | Early Telophase, Both POK1 and TAN1 at the division site | 100%, n = 14/14 | 39%, n = 12/31*** (p = 0.0001) |
|  | Early Telophase, Both POK1 and TAN1 at the division site, POK1 in the phragmoplast midline | 0%, n = 0/14 | 26%, n = 8/31* (p = 0.0436) |
|  | Early Telophase, NO POK1 or TAN1 at division site or phragmoplast midline | 0%, n = 0/14 | 35%, n = 11/31** (p = 0.098) |
|  | Late Telophase, Both POK1 and TAN1 at the division site only | 95%, n = 60/63 | 71%, n = 42/59*** (p = 0.0004) |
|  | Late Telophase, POK1 and TAN1 at the division site and POK1 in the phragmoplast midline | 5%, n = 3/63 | 19%, n = 11/59* (p = 0.022) |
|  | Late Telophase, TAN1 only at the division site | 0%, n = 0/63 | 7%, n = 4/59 (NS, p = 0.0518) |
|  | Late Telophase, Neither TAN1 or POK1 at division site or in the phragmoplast midline | 0%, n = 0/63 | 3%, n = 2/59 (NS, p = 0.2318) |

## Discussion

In *tan1* and *air9* single mutants, POK1 localizes to the division site and there are no discernable division plane defects (Model in Supplementary Figure 6). However, in the *tan1 air9* double mutant, POK1 co-localizes with the PPB but is lost from the division site during metaphase (Model in Figure 7). First, this suggests that TAN1 and AIR9 are not essential for POK1 co-localization with the PPB. Second, it suggests that POK1 is maintained at the division site after PPB disassembly via direct or indirect interactions with TAN1 or AIR9. We provide evidence that TAN1 interacts with POK1 through motifs within the first 132 amino acids of TAN1, as identified using the yeast-two-hybrid system. Alignments of TANGLED1 proteins from representative monocots and dicots, such as *Solanum lycopersium, Oryza sativa, Sorghum bicolor, Zea mays*, and *Brassica napus*, showed that amino acids 28-33 (INKVDK) are highly conserved across plant species (Supplementary Figure 7). Amino acids 30-32 (VDK) are identical and the remaining residues within the motif have similar properties across these plant species. The high degree of conservation suggests that these amino acids are likely important for TAN1 function. When alanine substitutions of these amino acids were introduced into TAN1 and transformed into Arabidopsis *tan1 air9* double mutants, we observed reduced TAN1 and POK1 localization at the division site, as well as defects in phragmoplast positioning. Here we hypothesize that amino acids 28-33 are essential for TAN1 and POK1 interaction in both yeast-two-hybrid and in Arabidopsis. In addition to several reports showing that TAN1 and POK1 interact using the yeast-two-hybrid system (Müller et al., 2006; Rasmussen et al., 2011), bimolecular fluorescence complementation has also been used to show TAN1-POK1 interactions in Arabidopsis protoplasts (Lipka et al., 2014). Alanine substitutions at positions 28-33 of TAN1 may disrupt TAN1-POK1 interaction through misfolding that blocks the POK1 interaction site or by affecting the amino acids that directly mediate POK1 binding. Regardless of the exact mechanism(s) of POK1-TAN1 physical interactions or the possibility that yeast-two-hybrid interactions do not reflect equivalent POK1-TAN1 physical interactions in Arabidopsis, we show that these TAN1 amino acids are involved in mediating TAN1 and POK1 localization to the division site.

**Figure 7:**
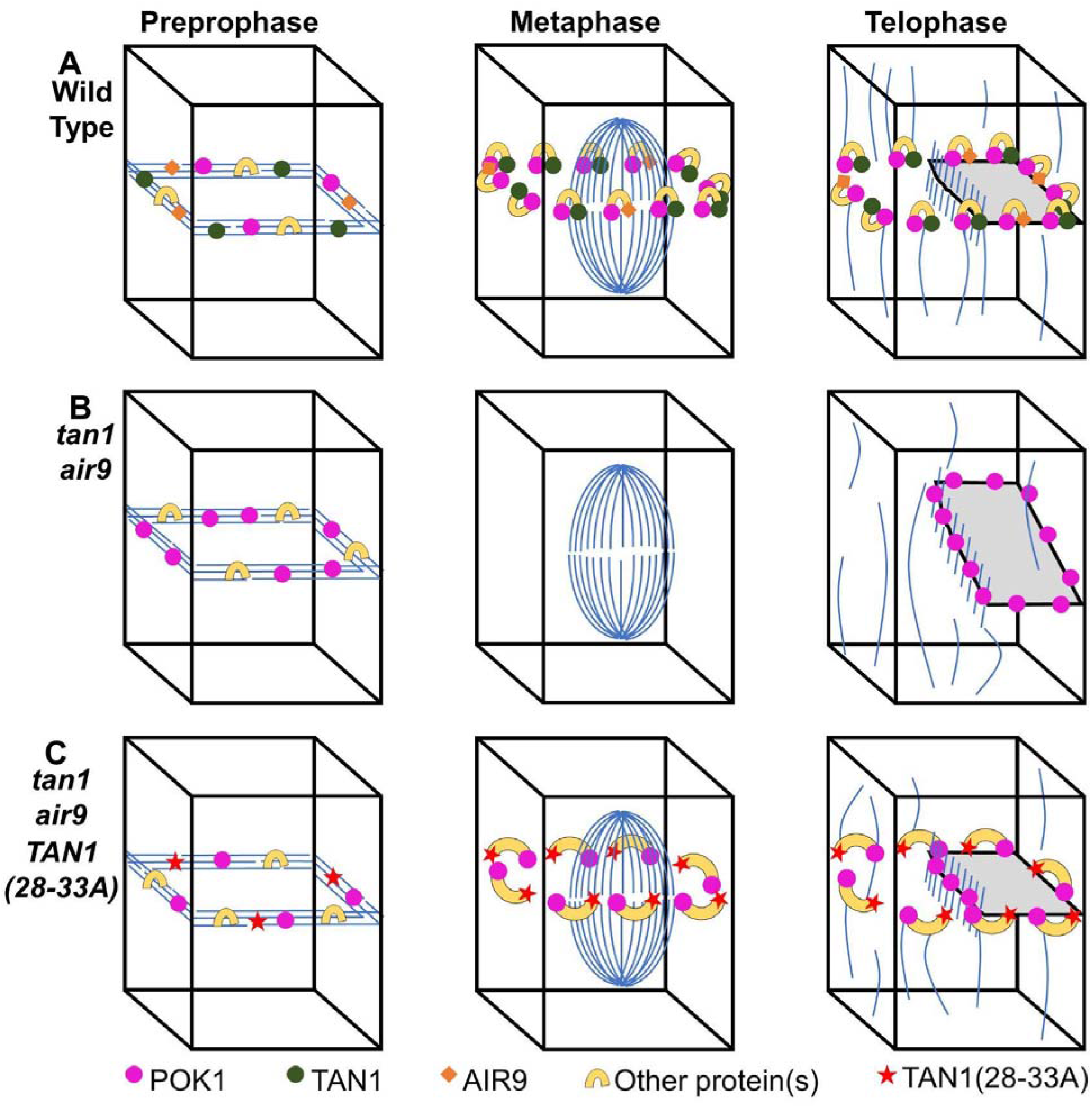
A speculative model on TAN1, AIR9, and POK1 interactions to ensure correct division plane orientation. A) In wild-type (WT) cells, AIR9, TAN1, and POK1 are recruited independently of one another to the PPB. Interaction between TAN1 and POK1 maintain both proteins at the division site through telophase, with AIR9 being re-recruited to the division site in late telophase. B) In the *tan1 air9* double mutant, TAN1, AIR9, and potential AIR9/POK1 interacting proteins are recruited to the PPB. Upon disassembly of the PPB, POK1 is lost from the division site and during telophase aberrantly accumulates in the phragmoplast midline. Due to the loss of TAN1 and POK1 from the division site, the phragmoplast is not guided to the location defined by the PPB. C) In *tan1 air9* double mutants expressing *TAN1(28-33A)*, TAN1(28-33A) and POK1 are recruited to the PPB independently of one another. POK1 and TAN1(28-33A) are partially maintained in some metaphase and early telophase cells possibly by interactions with other proteins. However, due to the inability of TAN1(28-33A) and POK1 to interact with one another, both proteins are not efficiently maintained at the division site. The majority (90%) of late telophase cells contain both POK1 and TAN1(28-33A) at the division site. Late recruitment of POK1 and TAN1(28-33A) may help guide the phragmoplast to the correct division site in a majority of cells.

We demonstrate that the first region of the TAN1 protein, the first 132 amino acids which primarily accumulates at the division site during telophase (Rasmussen et al., 2011), is both necessary (Mir et al., 2018) and sufficient to largely rescue the *tan1 air9* double mutant (Figure 2). This suggests that TAN1_1-132_ and its recruitment to the division site during telophase is critical for correct division plane orientation in the *tan1 air9* double mutant. Although full length TAN1 localizes to the division site throughout cell division, the ability of TAN1_1-132_ to rescue the *tan1 air9* double mutant, suggests that TAN1, and possibly POK1, localization to the PPB and division site during metaphase may not be required for division site maintenance in Arabidopsis. Indeed, whether the PPB itself is required for division plane positioning has been raised by analysis of a triple mutant in three closely related *TONNEAU RECRUITING MOTIF6,7,8* genes (*trm6,7,8). trm678* mutants, which lacked well defined PPBs, had disrupted POK1 recruitment to the division site but only minor defects in division positioning (Schaefer et al., 2017). However, when amino acids critical for TAN1-POK1 interactions in the yeast-two-hybrid system are disrupted by transforming *TAN1(28-33A)_1-132_-YFP* into the *tan1 air9* double mutant, root growth and phragmoplast positioning are disrupted. *TAN1(28-33A)_1-132_-YFP* accumulation at the division site during telophase was reduced compared to unaltered *TAN1_1-132_-YFP*. This suggests that TAN1-POK1 interaction promotes, but is not strictly necessary, for TAN1 recruitment to the division site during telophase.

Full-length TAN1 (*28-33A*) localizes to the division site throughout cell division and almost fully rescues the *tan1 air9* double mutant. TAN1, AIR9, and POK1 colocalize at the PPB independently of one another, which may promote the formation of protein complexes required for division site maintenance. Colocalizing with the PPB may provide an opportunity for proteins in close proximity to form stabilizing interactions before PPB disassembly. This suggests that recruitment of TAN1 and POK1 to the division site early in cell division may provide another temporally distinct way to promote correct division plane positioning. Phragmoplast positioning defects in TAN1(28-33A) *tan1 air9* plants may be the result of defects in phragmoplast guidance in cells that lacked TAN1(28-33A) and POK1 at the division site in metaphase or early telophase that were not corrected in late telophase.

The ability of TAN1(28-33A) and POK1 to remain at the division site in some cells after PPB disassembly in the *tan1 air9* double mutant suggests that there are other proteins that interact with TAN1 and/or POK1 that help stabilize them at the division site perhaps via the formation of multiprotein complexes. The pleckstrin homology GAPS, PHGAP1 and PHGAP2 (Stöckle et al., 2016), RANGAP1 (Xu et al., 2008), and IQ67 DOMAIN (IQD)6,7,8 proteins (Kumari et al., 2021) are division site localized proteins that may stabilize TAN1 and POK1 at the division site via their interaction with POK1. PHGAP1, PHGAP2, and RANGAP1 are dependent on POK1 and POK2 for division site recruitment. Like TAN1, RANGAP1 colocalizes with the PPB and remains at the division site throughout cell division (Xu et al., 2008). PHGAP1 and PHGAP2 are uniformly distributed in the cytoplasm and on the plasma membrane in interphase cells and accumulate at the division site during metaphase. These proteins also have their own distinct roles in division site maintenance (Stöckle et al., 2016). PHGAP2 has a likely role in division site establishment by regulating ROP activity (Hwang et al., 2008). RANGAP1 regulation of local RAN-GTP levels has potential roles in microtubule organization and division site identity (Xu et al., 2008). IQD6, IQD7, and IQD8 interact with POK1 and play a role in PPB formation and POK1 recruitment to the division site. *iqd678* triple mutants have PPB formation defects and fail to recruit POK1 to the division site in cells lacking PPBs. However, POK1 localization to the division site in *iqd678* mutants recovers during telophase to wild-type levels (Kumari et al., 2021). We speculate that this IQD6-8 independent recruitment may depend on TAN1. Unlike the PHGAPs and RANGAP1, IQD8 localization to the division site is not dependent on POK1 and POK2. This suggests that IQD6-8 proteins work upstream of POK1 to establish the division site and are important for POK1 recruitment to the division site early in cell division. Although TAN1-POK1 interaction becomes critical for TAN1 and POK1 maintenance at the division site in the absence of AIR9, other division site localized proteins may provide additional stability and help maintain TAN1 and POK1 at the division site.

How AIR9 stabilizes POK1 at the division site in the absence of TAN1 is less clear. There is no information about whether POK1 and AIR9 interact directly with one another. Additionally, AIR9 localization, in contrast to TAN1 localization, is intermittent at the division site. When expressed in tobacco Bright Yellow2 (BY2) cells, AIR9 colocalizes with the PPB but is then lost from the division site until late telophase when the phragmoplast contacts the cortex (Buschmann et al., 2006). In Arabidopsis, AIR9 may localize to the division site during metaphase or telophase, but it is difficult to observe because AIR9 also strongly colocalizes with cortical microtubules which may obscure AIR9 localization in nearby cells (Buschmann et al., 2015). Rather than directly interacting with POK1, AIR9 may recruit other proteins to the division site during preprophase that help maintain POK1 at the division site in the absence of TAN1. One potential candidate is the kinesin-like calmodulin binding protein, KCBP, which interacts with AIR9 (Buschmann et al., 2015). KCBP is a minus-end-directed kinesin (Song et al., 1997) that localizes to the division site in Arabidopsis and moss (Miki et al., 2014; Buschmann et al., 2015). We speculate that other TAN1, AIR9, and POK1 interacting proteins that have not been identified yet may be key for TAN1-POK1 division site maintenance.

POK1 and POK2 have roles in phragmoplast guidance, but POK2 also has a role in phragmoplast dynamics (Lipka et al., 2014; Herrmann et al., 2018). Although POK1 does not frequently accumulate in the phragmoplast midline in wild-type cells, POK2 showed striking dual localization to both the phragmoplast midline and the division site. Localization of POK2 to the phragmoplast midline required the N-terminal motor domain, while the C-terminal region was localized to the division site (Herrmann et al., 2018). Our hypothesis is that the “default” location of both POK1 and POK2 is at microtubule plus-ends at the phragmoplast midline, based on likely or confirmed plus-end directed motor activity (Chugh et al., 2018). Interactions with division site localized proteins, such as TAN1 and AIR9, may stabilize or recruit POK1 and POK2 at the division site away from the phragmoplast midline.

We demonstrate that AIR9 and TAN1 function redundantly to maintain POK1 at the division site to ensure correct cell wall placement. In the absence of AIR9, our data suggests that TAN1-POK1 interaction promotes, but is not required for, the maintenance of both proteins at the division site and disrupting this interaction partially disrupts their localization to the division site. This also suggests that other TAN1, POK1, and AIR9 interacting proteins are involved with stabilizing TAN1 and POK1 at the division site.

## Materials and methods

### Growth conditions, genotyping mutants, and root length measurements

Arabidopsis seedlings were grown on ½ strength Murashige and Skoog (MS) media (MP Biomedicals; Murashige and Skoog, 1962) containing 0.5 g/L MES (Fisher Scientific), pH 5.7, and 0.8% agar (Fisher Scientific). Seeds sown on plates were first stratified in the dark at 4°C for 2 to 5 days then grown vertically in a growth chamber (Percival) with 16/8-h light/dark cycles and temperature set to 22°C. For root length experiments, *tan1 air9* transgenic T3 lines expressing *p35S:TAN1-YFP, 35S:TAN1_1-132_-YFP, 35S:TAN1(28-33A)_1-132_-YFP*, or *35S:YFP-TAN1(28-33A)* were grown vertically, the plates were scanned (Epson) and root lengths were measured using FIJI (ImageJ, http://fiji.sc/) after 8 days. Untransformed *tan1 air9* double mutants and *air9* single mutants were grown alongside the double mutant seeds expressing the TAN1 constructs in equal numbers on the same plates to ensure plants were grown under the same conditions. After plates were scanned, seedlings were screened by confocal microscopy to identify seedlings expressing YFP translational fusion transgenes and CFP-TUBULIN, if present in the transgenic lines. At least 3 biological replicates, grown on separate plates on separate days, and at least 28 plants of each genotype total across all replicates were analyzed for each root growth experiment. Welch’s t-test was used to identify whether there were statistically significant differences between replicates before pooling the replicates for analysis. Root lengths were then plotted using Prism (GraphPad). Statistical analysis of root length was performed with Prism (GraphPad) using t-test with Welch’s correction. Welch’s t-test (unequal variance t-test) is used to test the hypothesis that two populations have equal means. Unlike the Student’s t-test, Welch’s t-test is often used when two samples have unequal variances or sample sizes. This test was used due to the unequal sample sizes because the plants examined were often segregating for multiple transgenes and had lower sample sizes than control plants such as *air9* single mutants and *tan1 air9* double mutants which either lacked transgenes or were segregating fewer transgenes.

YFP translational fusion TAN1 constructs were analyzed in *csh-tan (TAN1*, AT3G05330) *air9-31 (AIR9*, AT2G34680) double mutants in *Landsberg erecta (Ler*) unless otherwise specified. The *pPOK1:YFP-POK1* transgene in Columbia, a kind gift from Sabine Müller (Lipka et al., 2014), was crossed into the *tan-mad* and *air9-5* Columbia/Wassilewskija double mutant previously described (Mir et al., 2018). *tan-mad* and *air9-5* mutants were genotyped with primers ATRP and ATLP (to identify wild-type *TAN1*), JL202 and ATLP (to identify T-DNA insertion in *TAN1*), AIR9-5RP and AIR9-5LP (to identify wild-type *AIR9*), and LBb1.3 and AIR9RP (to identify T-DNA insertion in *AIR9*) and by observation of the *tan1 air9* double mutant phenotype (Supplementary Table 1).

### Generation of Transgenic Lines

*Agrobacterium tumefaciens*-mediated floral dip transformation was used as described (Clough and Bent, 1999). *csh-tan air9-31* double mutants were used for all floral dip transformations unless otherwise specified. Transgenic plants were selected on 15 μg/mL glufosinate (Finale; Bayer) and screened by microscopy before being transferred to soil and selfed. *CFP-TUBULIN* was crossed into *35S:TAN1(28-33A)_1-132_-YFP tan1 air9* plants using tan*1 air9 CFP-TUBULIN* plants (Mir et al., 2018) and progeny were subsequently screened by microscopy for CFP and YFP signal. *csh-tan1 air9-31* double mutants were confirmed by genotyping with primers ATLP and AtTAN 733-CDS Rw (to identify TAN1 wild-type), AtTAN 733-CDS Rw and Ds5-4 (to identify T-DNA insertion in *TAN1*), AIR9_cDNA 2230 F and AIR9 gnm7511 R (to identify *AIR9* wild-type), and AIR9 gnm7511 R and Ds5-4 (to identify T-DNA insertion in *AIR9*).

Columbia expressing the microtubule marker *UBQ10:mScarlet-MAP4* (Pan et al., 2020), a kind gift from Xue Pan and Zhenbiao Yang (UCR), was crossed to *tan-mad* and *air9-5* Columbia/Wassilewskija double mutants expressing *pPOK1:YFP-POK1*. Progeny were screened for mScarlet-MAP4 and YFP-POK1 by confocal microscopy and then selfed to recover *air9-5* single mutants, *tanmad* single mutants, and *air9-5 tan-mad* double mutants expressing *mScarlet-MAP4* and *YFP-POK1*.

*pTAN1:CFP-TAN1* and *pTAN1:CFP-TAN1(28-33A)* were introduced into *air9-5 tan-mad* double mutants expressing *mScarlet-MAP4* and *YFP-POK1* by *Agrobacterium tumefaciens*-mediated floral dip transformation. *pTAN1:CFP-TAN1* and *pTAN1:CFP-TAN1(28-33A)* transformants were selected on 100μg/mL gentamicin (Fisher Scientific) and the presence of mScarlet-MAP4, YFP-POK1, and either CFP-TAN1 or CFP-TAN1(28-33A) was confirmed by confocal microscopy. 4 independent transformed lines for *pTAN:CFP-TAN1(28-33A)* and 3 independent transformed lines for unaltered *pTAN:CFP-TAN1* were examined for division site localization cell counts.

### Plasmid Construction

*TAN1_1-132_-YFP* coding sequences were subcloned by EcoRI and BamHI double digestion from the plasmid *pEZRK-LNY-TAN1_1-132_-YFP* described previously (Rasmussen et al., 2011) into *pEZT-NL* vector (a kind gift from David Ehrhardt, Carnegie Institute, Stanford University) and selected with glufosinate (Finale; Bayer). The CFP-TUBULIN (CFP-TUA6) vector was previously described, a kind gift from Viktor Kirik (Kirik et al., 2007).

Six amino acid alanine substitutions were generated using overlapping PCR (primers in Supplementary Table 1) beginning at amino acid 10 of TAN1_1-132_. *TAN1_1-132_-YFP* coding sequence from plasmids described previously was used as the PCR template (Rasmussen et al., 2011). *TAN1(28-33A)_1-132_-YFP* was subcloned by EcoRI BamHI double digestion into pEZT-NL. To generate *YFP-TAN1(28-33A)*, alanine substitutions were first introduced into G22672 (TAN1 cDNA in pENTR223, from the Arabidopsis Biological Resource Center) using overlapping PCR with the same primers to generate *TAN1(28-33A)*. Gateway LR reaction (Fisher Scientific) was then used to subclone *TAN1(28-33A)* into pEarley104 (Earley et al., 2006).

*pTAN:CFP-TAN1(28-33A)* was generated using overlapping PCR. The TANGLED1 native promoter was amplified from *Np:AtTAN-YFP (Walker et al., 2007*) using the primers NpTANSacIFor and NpTANceruleanRev. Cerulean was amplified from the Cerulean CDS in pDONR221P4r/P3r using the primers NpTANceruleanFor and CeruleanpEarleyRev. TAN1(28-33A) in pEarley104 was amplified using CeruleanpEarleyFor and pEarleyOCSPstIRev. TANGLED1 native promoter, Cerulean, and TAN1(28-33A) were then combined using overlapping PCR using NpTANSacI and pEarleyOCSPstIRev. *pTAN:CFP-TAN1(28-33A)* was then subcloned into pJHA212G, a kind gift of Meng Chen (UCR), using SacI and PstI double digest. pTAN:CFP-TAN1 was generated the same way as pTAN:CFP-TAN1(28-33A) except unaltered TAN1 in pEarley104 was amplified using CeruleanpEarleyFor and pEarleyOCSPstIRev.

### Microscopy

An inverted Ti Eclipse (Nikon) with motorized stage (ASI Piezo) and spinning-disk confocal microscope (Yokogawa W1) built by Solamere Technology was used with Micromanager software (micromanager.org). Solid-state lasers (Obis) and emission filters (Chroma Technology) were used. For CFP translational fusions excitation 445, emission 480/40 was used; YFP translational fusions excitation 514, emission 540/30; and propidium iodide (PI), Atexa-568 goat anti-mouse antibody, and mScarlet-MAP4 excitation 561, emission 620/60 were used. The 20x objective has 0.75 numerical aperture and the 60x objective has 1.2 numerical aperture which was used with perfluorocarbon immersion liquid (RIAAA-6788 Cargille). Excitation spectra for mScarlet-MAP4 and YFP-POK1 partially overlapped, which resulted in faint bleed through signal in the YFP channel for some dense microtubule structures (e.g. spindles and phragmoplasts). YFP-POK1 colocalization with PPBs was carefully determined based on distinct YFP-POK1 signal and the presence of cytosolic YFP-POK1.

The ratio of the division site versus cytosolic fluorescence intensity was determined by taking the median YFP fluorescence intensity from the center Z-stack of individual cells with PPBs or phragmoplasts. For each cell the median fluorescence intensity was measured for two cytosolic areas and the division site on each side of the cell using circles with areas of 0.875 μm^2^. The sum of the median intensity at the division site on each side was then divided by the sum of the median intensity of the two cytosolic areas to calculate the ratio of the division site versus cytosolic fluorescence intensity. Fluorescence intensities were measured in FIJI. All plants used for this analysis were grown on the same day and imaged using identical conditions, and at least 5 plants of each genotype were examined.

### Measurements of PPB and phragmoplast angles and cell file rotation

At least 3 biological replicates, grown on separate plates on separate days, composed of at least 15 plants per genotype for PPB measurements and at least 8 plants per genotype for phragmoplast measurements were used to gather angle data. 8-day-old seedlings were stained with 10 μM PI for 1 minute and then destained in distilled water before imaging by confocal microscopy using a 20x or 60x objective. PPB and phragmoplast angles were measured using FIJI. The angle was measured between the left-hand cell wall and the orientation of the PPB or phragmoplast in the root tips of *tan1 air9* double mutants expressing *CFP-TUBULIN* or immunostained microtubules (described in the next section). Cell file rotation was examined by measuring from the left-hand side of the transverse cell wall relative to the long axis of the root in images of the differentiation zone stained with PI. The differentiation zone was identified by the presence of root hairs. Prism (GraphPad) and Excel (Microsoft Office) were used to perform statistical analyses and to plot data. F-test was used to compare normally distributed variances (PPB and phragmoplast angles) and Levene’s test was used to compare non-normally distributed variances (cell file rotation angle measurements). *tan1 air9* double mutants have non-normally distributed cell file twisting because the roots tend to twist to the left (Mir et al., 2018). Genotypes across biological replicates were compared to ensure there was no statistically significant difference between them before pooling data.

### Immunostaining

*air9, tan1 air9 p35S:TAN1-YFP, tan1 air9 35S:YFP-TAN1(28-33A)*, and untransformed *tan1 air9* plants were stratified and then grown vertically on ½ MS plates in a growth chamber at 22°C with a 16/8-h light/dark cycle for 8 days. The seedlings were screened by microscopy for YFP and then fixed and processed for immunofluorescence microscopy using a 1:2000 dilution of monoclonal anti-α-tubulin B-5-1-2 antibody (Life Technologies; 32-2500) followed by 1:2000 dilution of Alexa-568 goat anti-mouse antibody (Thermo Fisher; A-11004) as described previously (Sugimoto et al., 2000).

### Yeast-two-hybrid

Six alanine substitutions were generated using overlapping PCR and TAN1 coding sequence in pEZRK-LNY-*TAN1_1-132_-YFP* as a template beginning at amino acid 10 of TAN1 and continuing through amino acid 123 according to (Russell and Sambrook, 2001). All except amino acids substitutions for 64-69 and 106-111 were cloned into pAS vector (Fan et al., 1997) using EcoRI BamHI double digestion. pBD-TAN1(28-33A) was generated by using primers Ala_05_FOR and Ala_05_REV to perform DpnI mediated site-directed mutagenesis by PCR (Fisher and Pei, 1997). pBD-TAN1 (Walker et al., 2007), and *pAS-TAN1_1-132_* (Rasmussen et al., 2011) were used as positive controls, while pAD-MUT was used as a negative control for testing interaction with pAD-POK1 (Müller et al., 2006). *pAD-POK1* and *pAS-TAN1_1-132_* constructs were co-transformed into yeast strain YRG2 according to manufacturer instructions (Stragene). Positive yeast-two-hybrid interaction was determined by the presence of growth on plates cultured at 30°C lacking histidine after 3 days. Plates were then scanned (Epson).

## Acknowledgements

Thanks to Andrew Gomez (UCR, supported by USDA-NIFA 2017-38422-27135) for help with yeast-two-hybrid experiments, Prof. Sabine Müller (University of Tübingen) for YFP-POK1 seeds, and Profs. Meng Chen and David Nelson (UCR) for their helpful comments on alanine scanning mutagenesis. Thanks to Prof. Henrik Buschmann (Osnabrück University) for original *tan1 air9* characterization. Thanks to Lindy Allsman, Stephanie Martinez, and Aimee Uyehara (UCR) for helpful comments on the manuscript. NSF-CAREER #1942734 and NSF-MCB #1716972, USDA-NIFA-CA-R-BPS-5108-H are gratefully acknowledged for funding.

## Supplementary Figures

**Supplementary Figure 1.**
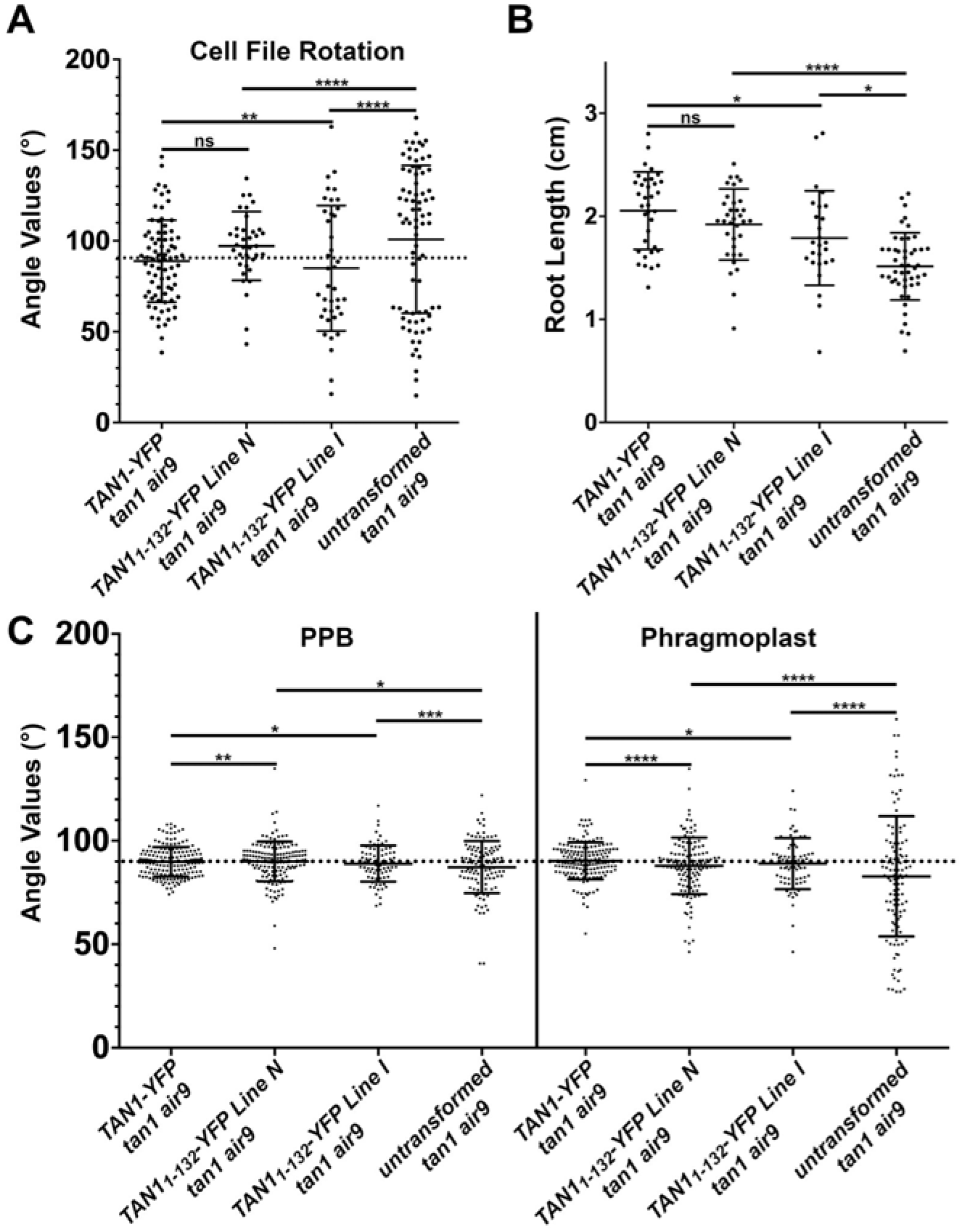
*p35S:TAN1_1-132_-YFP tan1 air9* lines show significant rescue compared to untransformed *tan1 air9* double mutants. A) Cell file rotation angles of *tan1 air9* double mutants expressing *p35S:TAN1-YFP* (left), two *p35S:TAN1_1-132_-YFP* transgenic lines designated as line N (center left) and line I (center right) and untransformed *tan1 air9* plants (right) n > 6 plants for each genotype. Angle variances were compared with Levene’s test. B) Root length measurements from 8 days after stratification of *tan1 air9* double mutants expressing *p35S:TAN1-YFP* (left), two *p35S:TAN1_1-132_-YFP* transgenic lines (middle) and untransformed plants (right), n > 13 plants for each genotype, compared by two-tailed t-test with Welch’s correction. C) PPB and phragmoplast angle measurements in dividing root cells of *tan1 air9* double mutants expressing *p35S:TAN1-YFP* (left), two *p35S:TAN1_1-132_-YFP* transgenic lines (middle) and untransformed plants (right), n > 23 plants of each genotype. Angle variations compared with F-test ns indicates not significant, * P-value <0.05, ** P-value <0.01, **** P-value <0.0001.

**Supplementary Figure 2.**
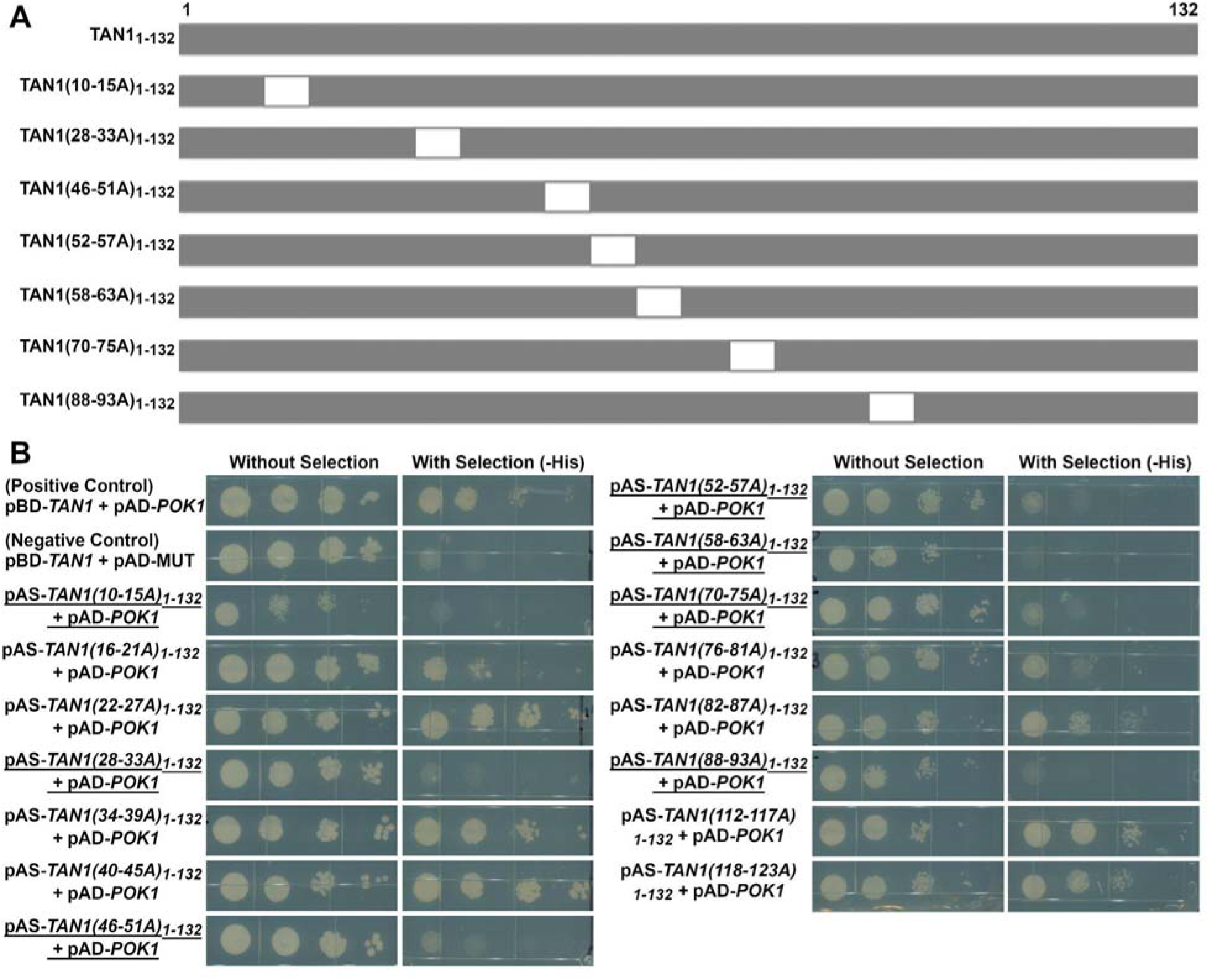
Yeast-two-hybrid interactions between POK1 (C-terminal amino acids 1683-2066, as previously described (Müller et al., 2006; Rasmussen et al., 2011; Lipka et al., 2014)) and TAN1_1-132_ alanine scanning constructs. A) Diagram of alanine scanning constructs that showed loss of interaction with POK1 by yeast-two-hybrid. The location of the six alanine substitutions within each TAN1_1-132_ construct are represented by white boxes. B) Yeast-two-hybrid results of screen for loss of interaction with POK1. Underlined constructs showed loss of interaction with POK1. Alanines 64-69 and 106-111 were not completed and not included in the yeast-two-hybrid.

**Supplementary Figure 3.**
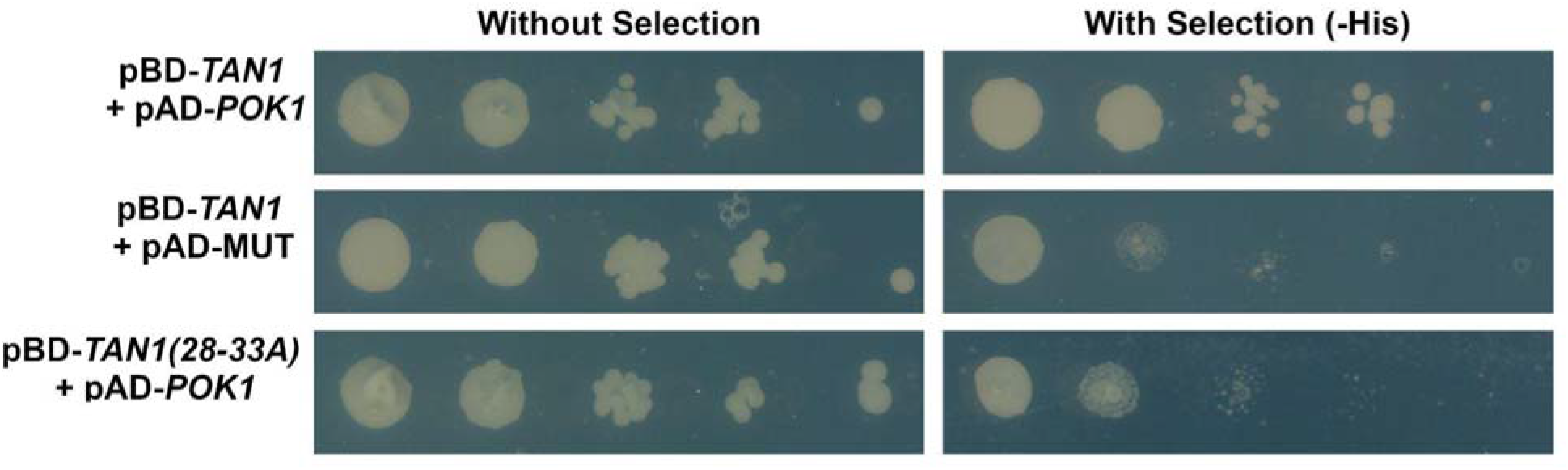
Yeast-two-hybrid interactions between TAN1 and POK1 (C-terminal amino acids 1683-2066, as previously described (Müller et al., 2006; Rasmussen et al., 2011; Lipka et al., 2014)) and TAN1(28-33A). Full-length TAN1(28-33A) does not interact with POK1 by yeast-two-hybrid.

**Supplementary Figure 4.**
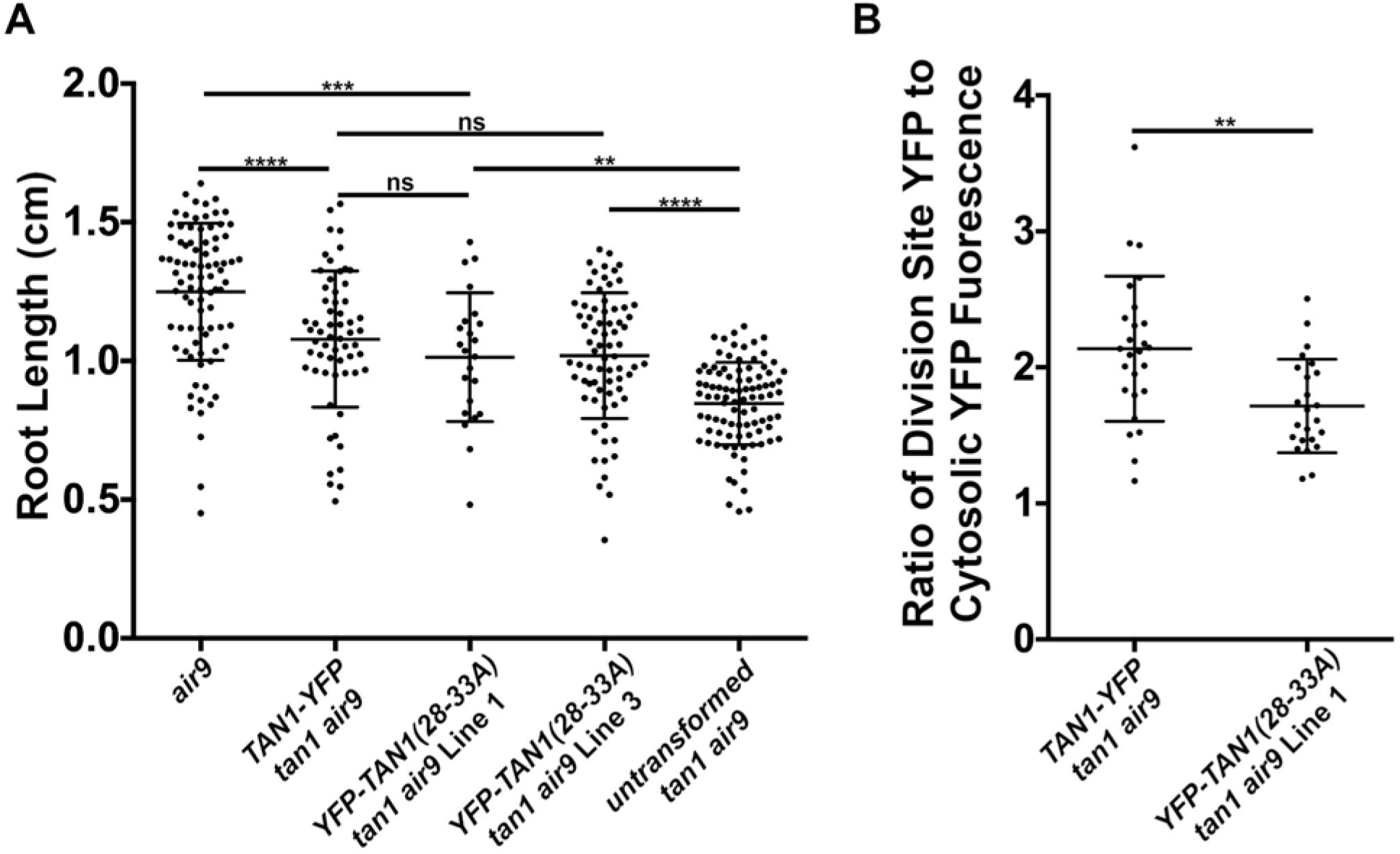
*p35S:YFP-TAN1(28-33A) tan1 air9* lines show significant rescue compared to untransformed *tan1 air9*, but less accumulation of YFP-TAN1(28-33A) during telophase. A) Root length measurements from 8 days after stratification of *air9* single mutants (left), tan*1 air9* double mutants expressing *p35S:TAN1-YFP* (second from the left), two *p35S:YFP-TAN1(28-33A)-YFP* transgenic lines designated as line 1 (center) and line 3 (second from the right), and untransformed plants (right), n > 22 plants for each genotype, compared by two-tailed t-test with Welch’s correction. B) Ratio of TAN1-YFP or TAN1(28-33A)-YFP fluorescence at the division site to cytosolic fluorescence from *tan1 air9* plants expressing *p35S:TAN1-YFP* (left) or *p35S:YFP-TAN1(28-33A)* (right) during telophase, n >12 plants for each genotype. Asterisks indicate a significant difference as determined by Mann-Whitney U test. ns indicates not significant, ** P-value <0.01, *** P-value <0.001, **** P-value <0.0001. Note: TAN1-YFP fluorescence measurements are the same as those used for the telophase fluorescence measurements in supplementary figure 5E.

**Supplementary Figure 5.**
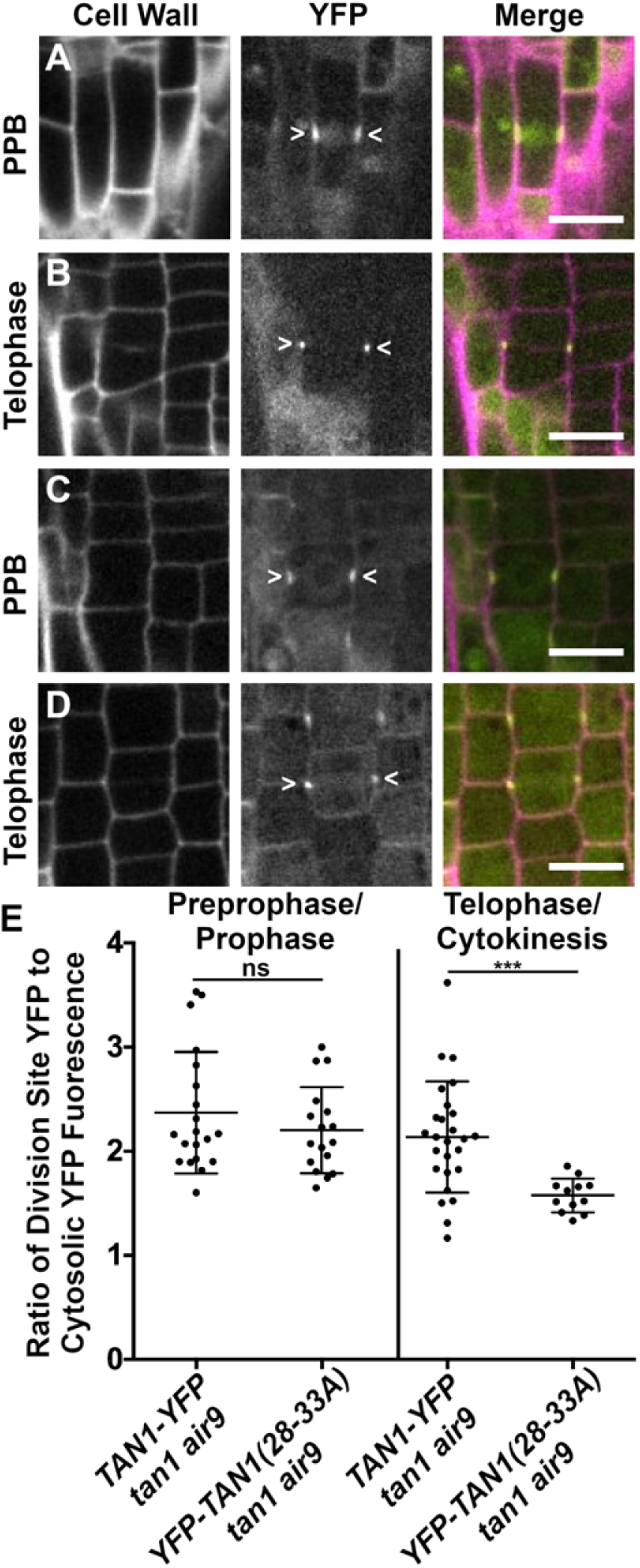
YFP-TAN1(28-33A) localizes to the division site in preprophase or prophase and with reduced fluorescence during telophase in *tan1 air9* mutants. Propidium iodide stained *tan1 air9* plants expressing *p35S:TAN1-YFP* in (A) preprophase or prophase (B) telophase or cytokinesis. *tan1 air9* plants expressing *p35S:YFP-TAN1(28-33A)* in (C) preprophase or prophase (D) telophase or cytokinesis. The division site is indicated by arrowheads in the YFP panels. Bars = 10 μm. E) TAN1-YFP or TAN1(28-33A)-YFP ratio of the division site versus cytosolic fluorescence intensity from *tan1 air9* plants expressing *p35S:TAN1-YFP* or *p35S:YFP-TAN1(28-33A)* during preprophase or prophase and telophase or cytokinesis, n >5 plants for each genotype. Ratios compared with Mann-Whitney U test. ns indicates not significant, *** P-value <0.001.

**Supplementary Figure 6.**
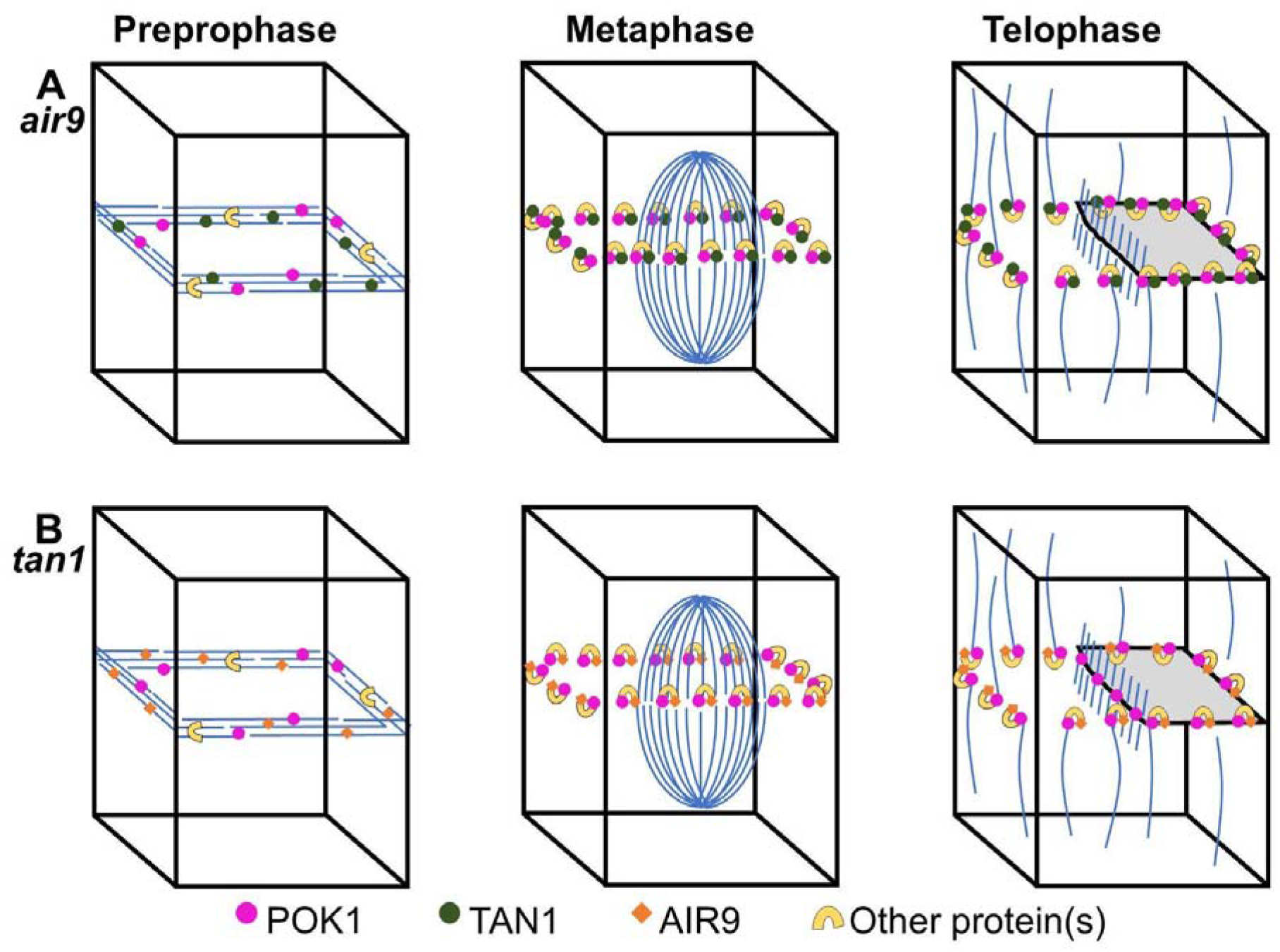
A model of POK1 localization in *tan1* and *air9* single mutants. A) In *air9* single mutants TAN1 and POK1 are recruited to the PPB and their interaction with one another and other proteins stabilizes TAN1 and POK1 at the division site. B) In *tan1* single mutants AIR9 and POK1 are recruited to the PPB. POK1 is potentially stabilized at the division site either by interacting directly with AIR9 or another protein recruited to the division site by AIR9. POK1 tends to accumulate in the phragmoplast midline in *tan1* single mutants, which may reflect that POK1 is not as efficiently recruited to the division site in the absence of TAN1.

**Supplementary Figure 7.**
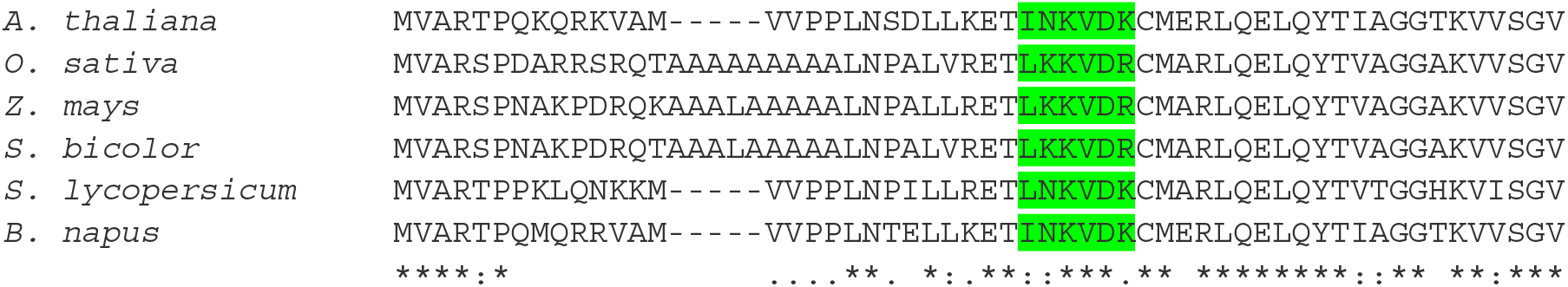
Alignments of amino acids 1-55 of *A. thaliana* TAN1 with TAN1 homologs from other plant species. Amino acids 28-33 of Arabidopsis TAN1 and amino acids that align with them in other plant species are highlighted in green. “*” indicates residues are fully conserved, “:” indicates strong conservation of properties across species, and “.” indicates weak conservation of properties across species.

**Supplementary Table 1.**
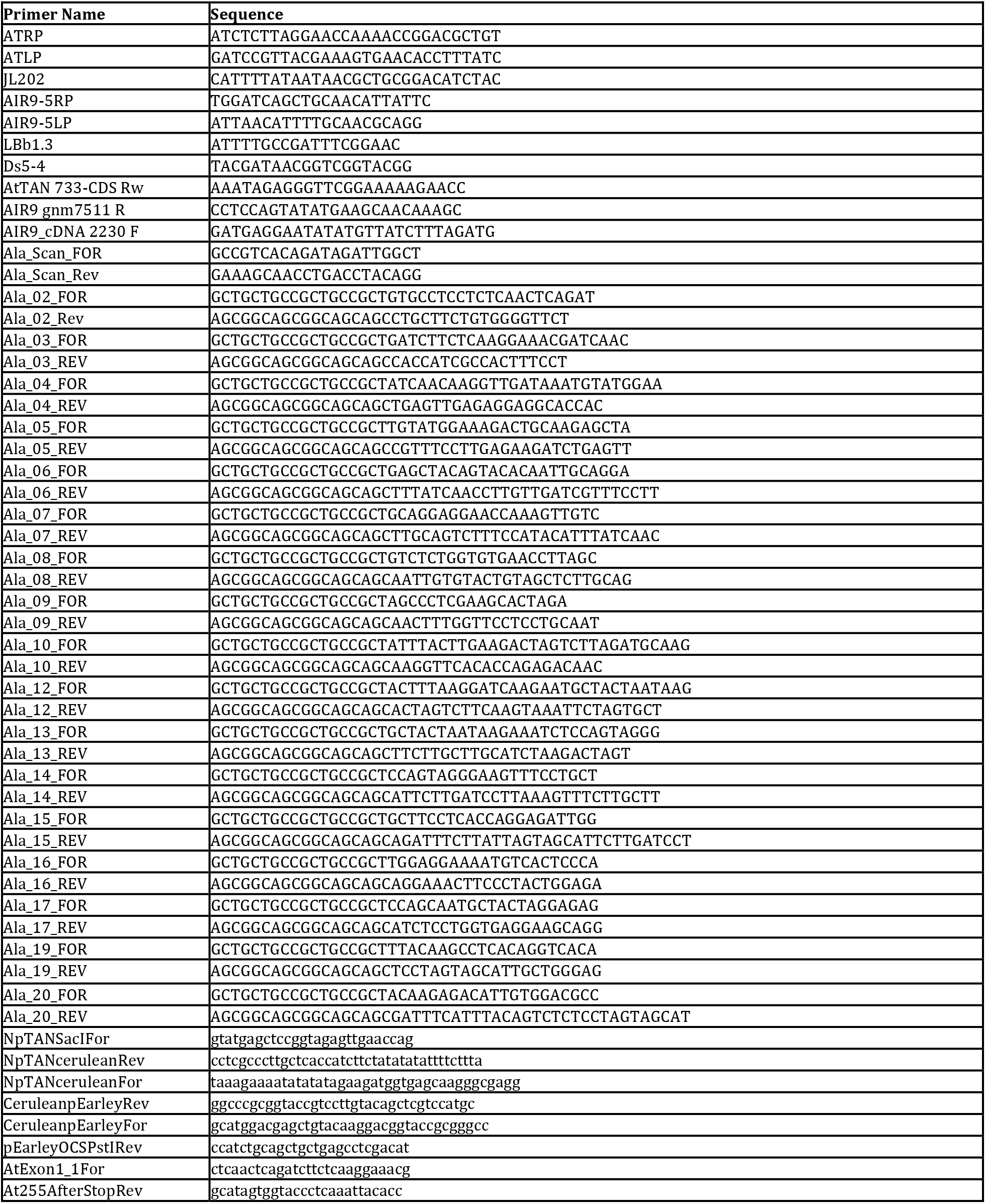
Primers used for cloning and genotyping.

